# EFCAB9 is a pH-Dependent Ca^2+^ Sensor that Regulates CatSper Channel Activity and Sperm Motility

**DOI:** 10.1101/459487

**Authors:** Jae Yeon Hwang, Mannowetz Nadja, Yongdeng Zhang, Robert A. Everley, Steven P. Gygi, Joerg Bewersdorf, Polina V. Lishko, Jean-Ju Chung

## Abstract

**Summary:** Varying pH of luminal fluid along the female reproductive tract is a physiological cue that modulates sperm motility. CatSper is a sperm-specific, pH-sensitive calcium channel essential for hyperactivated motility and male fertility. Multi-subunit CatSper channel complexes organize linear Ca^2+^ signaling nanodomains along the sperm tail. Here, we identify EF-hand calciumbinding domain-containing protein 9 (EFCAB9) as a dual function, cytoplasmic machine modulating the channel activity and the domain organization of CatSper. Knockout mice studies demonstrate that EFCAB9, in complex with the CatSper subunit, CAT SPERζ, is essential for pH-dependent and Ca^2+^ sensitive activation of the CatSper channel. In the absence of EFCAB9, sperm motility and fertility is compromised and the linear arrangement of the Ca^2+^ signaling domains is disrupted. EFCAB9 interacts directly with CATSPERζ in a Ca^2+^ dependent manner and dissociates at elevated pH.These observations suggest that EFCAB9 is a long-sought, intracellular, pH-dependent Ca^2+^ sensor that triggers changes in sperm motility.

**Highlights:**

- *Efcab9* encodes an evolutionarily conserved, sperm-specific EF-hand domain protein
- *Efcab9*-deficient mice have sperm motility defects and reduced male fertility
- EFCAB9 is a pH-tuned Ca^2+^ sensor for flagellar CatSper Ca^2+^ channel
- EFCAB9 is a dual function machine in gatekeeping and domain organization of CatSper

## INTRODUCTION

Changes in sperm motility patterns (Yanagimachi, 1970; Yanagimachi, 2017), observed in the steering chemotactic movement of marine invertebrate spermatozoa (Bohmer et al., 2005; Wood et al., 2005) and triggering hyperactivated motility in mammals (Ho and Suarez, 2001; Suarez et al., 1993 require Ca^2+^ influx across the flagellar membrane. The molecular commonalities for this Ca^2+^ entry pathway into these sperm cells are the flagella-specific and Ca^2+^-selective channel,(CarlSon et al., 2003; Qi et al., 2007; Quill et al., 2003; Ren et al., 2001; Seifert et al.,2015) and its activation by intracellular alkalinization (Kirichok et al., 2006; Lishko et al., 2010; Lishko et al., 2011; Miller et al., 2016; Seifert et al., 2015; Strunker et al., 2011).This implies that an evolutionarily conserved pH- and Ca^2+^-sensing mechanism regulates the CatSper channel.

The CatSper channel is the most complex of all known ion channels, encoded by at least nine genes in mammals. It is comprised of pore-forming α subunits (CatSper 1-4) (Qi et al., 2007; Quill et al., 2003; Ren et al., 2001) and five accessory subunits (transmembrane CatSper α, γ, δ,ε, and cytosolic CatSper) (chung et al., 2017; Chung et al., 2011; Liu et al., 2007; Wang et al., 2009. Male mice lacking CatSper genes (Chung et al., 2017; Chung et al., 2011; Quill et al., 2003; Ren et al., 2001), as well as humans with loss of function mutations (Avenarius et al., 2009; Hildebrand et al., 2010; Smith et al., 2013) in transmembrane subunits, are completely infertile due to lack of sperm hyperactivation. More mechanistic understanding of the regulatory mechanisms has been hampered by the inability to heterologously reconstitute the complex channel and the inseparability of phenotypes resulting from loss of each subunit in sperm cells (Chung et al., 2017; Chung et al., 2011). Thus, the molecular mechanism of this pH-dependent activation of CatSper has relied on speculation, based on amino acid properties and sequence homology of the subunits. Moreover, to date, none of the known CatSper subunits contained calcium /CaM binding motifs. This suggests that additional proteins assist complex assembly and trafficking, and/or detect changes in intracellular pH and Ca^2+^ for the CatSper channel.

The recent identification of CatSperζ as a unique cytoplasmic component of the mammalian CatSper channel complex has been illuminating. It was previously reported that CatSper organizes Ca^2+^ signaling nanodomains, uniquely aligned along the sperm tail as ‘racing stripes’ to form a network of intracellular signaling molecules (Chung et al.,2014). I n contrast to other CatSper knockout models, deficiency of CatSperζ disrupts the CatSper domains but the channel is functional and only results in *CatSperz*-null male subfertility (Chung et al., 2017; Chung et al., 2014). These studies indicate that CatSperζ functions in the compartmentalization of Ca^2+^signaling in mammalian sperm, and thus may modulate the Cainflux mechanism of the CatSper channel.

Here we use comparative proteomic and genomic screens and identify EFCAB9, a sperm-specific EF-hand protein, as a new auxiliary protein of the CatSper channel that modulates channel activity regulation and domain organization. We develop an *in vivo* mouse model to examine EFCAB9 function at the molecular, cellular, biochemical, electrophysiological, and super-resolution imaging levels. We find that EFCAB9 is a pH-dependent Ca^2+^ sensor for the CatSper channel. EFCAB9-CatSperζ forms a binary complex and interacts with the pore as a gatekeeper, activating CatSper channel responding to capacitation-associated intracellular changes. EFCAB9 interaction with CatSperζ requires its Ca^2+^ binding to EF-hands, which are responsive to pH changes in a physiologically relevant range. Loss of EFCAB9-CatSperζ largely eliminates the pH-dependent and Ca^2+^ sensitive activation of CatSper and alters newly resolved CatSper Ca^2+^ signaling domains. In contrast to the recent addition of CatSperζ to mammals, the EFCAB9 and other transmembrane CatSper subunits are evolutionarily conserved from flagellated single cell eukaryotes. These findings provide insight into evolutionarily conserved gating mechanisms for CatSper channels and reveal an adaptation for Ca^2+^ signaling in flagella.

## RESULTS

### Comparative Proteomic and Genomic Screens Identify Unknown Components of CatSper Calcium Channel Complex in Sperm

Sperm hyperactivated motility requires pH-dependent activation of the CatSper flagellar Ca^2+^ channel. The CatSper channel forms a multi-protein complex composed of at least nine subunits (CatSper1-4, β, γ, δ, ε, and ζ) (Chung at al.,2017). Here we examine a new member of the complex and determine its functional significance.

Our previous studies showed that other CatSper subunits were not detected in *CatSper1*-and *d*-null sperm cells despite their expression during spermatogenesis (Chung et al., 2017; Chung et al., 2011). To take advantage of this interdependence and identify new CatSper components, we performed a quantitative whole sperm proteomic screen to compare *CatSper1*-null with wild-type (wt) sperm (figure1 A), concomitant with our previous phosphotyrosine proteome analysis (Chung et al., 2014). Using tandem mass tag (TMT) labeling and mass spectrometry analysis, we quantified 3,227 proteins from *CatSper1*-null and wt spermatozoa, which is comparable to the size of other mouse sperm proteome (Castaneda et al., 2017). To identify the most significant differences of individual protein expression in biological replicates, we combined a measure of statistical significance with the magnitude of change to plot replicate data points(Figure 1A). In the experiment run in triplicate, only 19 proteins (14 downregulated and 5 upregulated significantly at *p*<0.05) were differentially expressed in *CatSper1*-null sperm by more than 2-fold (Figure 1A and Table S1) down from 73 candidate proteins (25 downregulated and 48 upregulated). This overall lack of differential protein expression between wt and *CatSper1*-null spermatozoa is particularly remarkable since our screen still identified all the nine known transmembrane and cytosolic components of the CatSper channel complex in a precise, quantitative, and reproducible manner (Figure 1A). This result validates the accuracy of the data and strongly suggests that mature *CatSper1*-null spermatozoa lack only proteins that normally form a stable complex with the CatSper channel. Thus, five additional downregulated proteins identified by our screen are strong candidates for CatSper components in spermatozoa that are unincorporated into the flagellar membrane or degraded in the absence of the CatSper channel.

**Figure 1.**
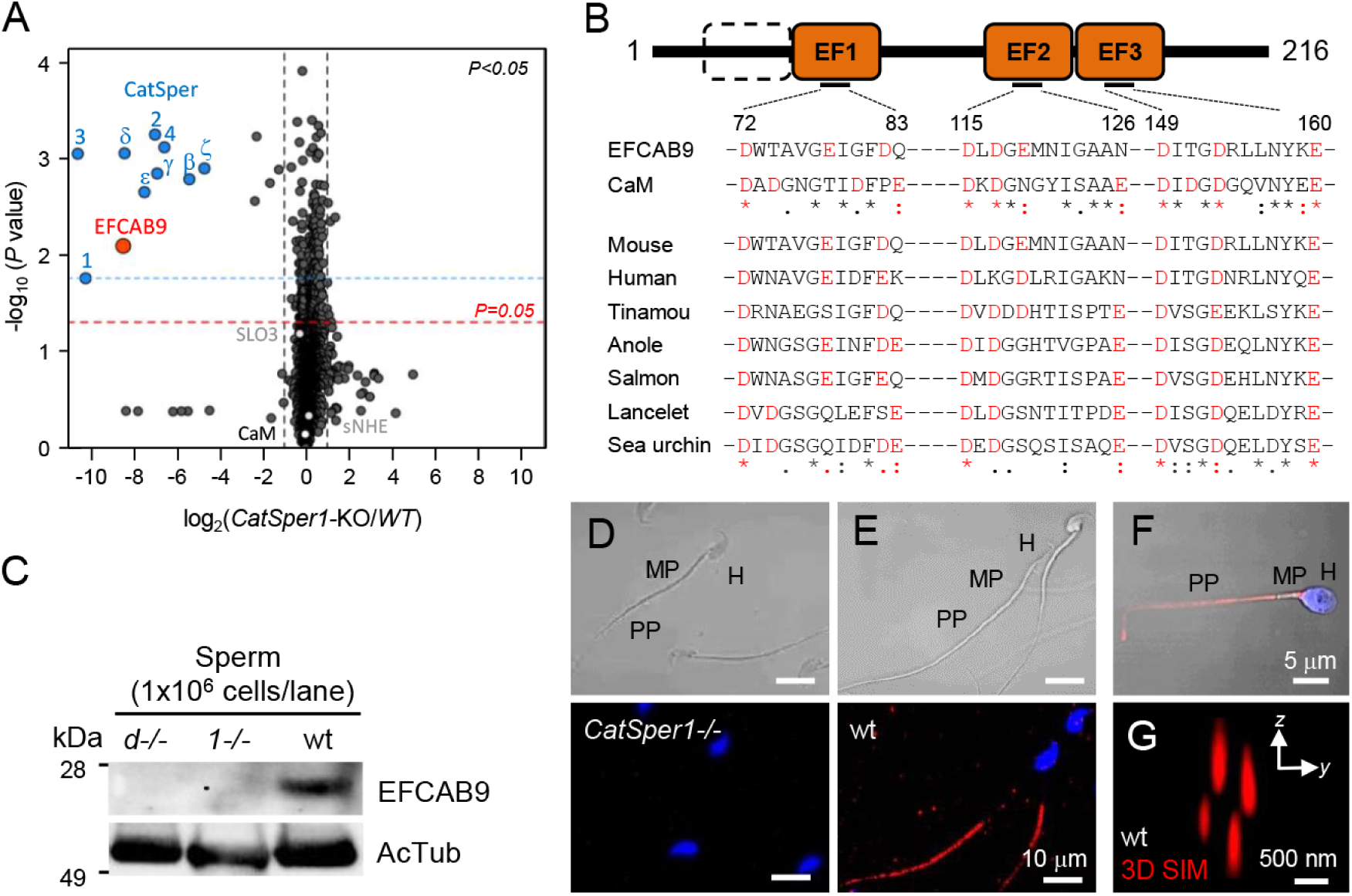
EFCAB9 is an intracellular protein associated with CatSper channel complex. (A) Quantitative analysis of whole sperm proteome from wt compared with *CatSper1*-/- mice to identify proteins associated with CatSper channel complex. Volcano plot of statistical significance against protein fold-change in *CatSper1*-/- over wt spermatozoa. Each protein is represented as a dot and is mapped according to its average fold change on the *x* axis and t-test p-value on the *y* axis. EFCAB9 (red dot) is one of the proteins with the most reduced protein expression in *CatSper1*-/- spermatozoa, clustered together with other CatSper subunits (blue dots) that are previously reported to express interdependently with CatSper1 in sperm cells. (B) The predicted domain structure of mouse EFCAB9 protein. Mouse EFCAB9 contains 216 amino acids with three predicted EF-hand domains represented with filled box. Empty box indicates the counterpart region aligned to the first EF-hand domain of calmodulin. Conserved acidic amino acids in the loop regions of EF-hand domains are highlighted in red. (C-E) Validation of EFCAB9 protein expression in sperm cells of wt mice compared with homozygous *CatSper1-* and CatSperd-null mice by immunoblot of total sperm extract (C) and immunofluorescence confocal images of mouse sperm (D and E). (F) Confocal immunofluorescent detection of EFCAB9 protein in human sperm. Red and blue signals indicate immunostained EFCAB9 and Hoechst-stained DNA, respectively (D-F). The corresponding DIC are shown for D and E. (G) A cross-section 3D SIM image of EFCAB9 in wt mouse sperm. *See also Figures S1 and S2*.

To further validate the candidates, we defined two additional criteria, based on the characteristics common to the known CatSper components: (1) be testis-specific and post-meiotically expressed, and (2) share evolutionary history with typical conservation patterns of lineagespecific gain and loss of *CatSper* genes. *Efcab9* meets both criteria of this combinatorial analyses of the transcriptome database of tissue expression (Yue et al.,2014)(Figure S1A) and comparative genomic screens (Figure S1B).

### EFCAB9I Is an Intraflagellar Protein with Conserved EF-Hand Ca^2+^ Binding Domains

EFCAB9 has three conserved EF-hand Ca^2+^ binding domains (Figure 1B). Amino acid sequence alignment of EFCAB9 to calmodulin (CaM) predicts that three conserved helix-loop-helix folding motifs (EF1-3) contain candidate EF-hand Ca^2+^ binding sites (Figure S2A and S2B). EF-hand Ca^2+^ binding sites are the side chains of aspartic and glutamic acids located in each loop (Figures 1B and S2A-S2C). Interestingly, sequence comparison of the EF loops of EFCAB9 orthologues revealed species-specific variations in EF1 and EF2 from the canonical EF loop containing an acidic residue at position 12 (Figure S2C). EFCAB9 was tightly clustered with all the other previously reported CatSper subunits as one of the proteins with the most reduced expression in *CatSper1-null* spermatozoa (Figure 1A and Table S1). We confirmed that *Efcab9* is expressed only in the testis, which contrasts with the genes encoding other EF-hand containing proteins quantified in our sperm proteomes including CaM and EFCAB1 (Figure S1C and TableS1). Interestingly, *Efcab9* transcripts are detected post-meiotically, similar to the genes encoding the CatSper pore-forming α-subunits, and unlike all the other CatSper auxiliary subunits (e.g.*CatSperz*, Figure S1D Chung et al., 2017; Chung et al., 2011).We compared EFCAB9 protein expression and subcellular localization in wt sperm cells with those in *CatSper1*-and *CatSperd*-null spermatozoa (Figures 1C-1E) by generating a specific antibody recognizing EFCAB9 (Figures S2D, S2E and S3C). EFCAB9 is localized to the principal piece of the wt mouse and human sperm tail (Figures 1E and 1F), as are other CatSper subunits, but is not detected in mouse sperm lacking the CatSper channel (Figure 1D). 3D structured illumination microscopy (SIM) imaging revealed that EFCAB9 also localizes in the characteristic four linear domains of sperm flagella (Figure 1G). From these results, we hypothesized that EFCAB9 is a functionally relevant CatSper channel component. The EF-hand is one of the major Ca^2+^-binding regions found on many of the Ca^2+^ sensors including CaM (Clapham, 2007 and EFCAB9 is the first CatSper component with a known, conserved domain. This raises the question of whether EFCAB9 underlies Ca^2+^ sensing of the CatSper channel in regulating sperm hyperactivated motility.

### EFCAB9-Deficient Mice Have Reduced Male Fertility and Phenotypes Resembling Loss of CatSperζ Function

To examine the potential role of EFCAB9 in the CatSper-mediated Ca^2+^ signaling pathway, we created *Efcab9* knockout mice by CRISPR/Cas9 genome editing (Figures 2 and S3). In our targeting strategy, gRNA was selected to introduce indels into the first exon of the *Efcab9* gene located on chromosome 11 (Figures S3A and S3B). We obtained two mutant alleles with either a 5 bp (*Efcab9-5del*) or a 28 bp (*Efcab9-28del*) deletion. These two mutant lines resulted in identical phenotypes in our initial characterization of bi-allelic homozygous (null) mice, ruling out the possibility that off-target effects account for the phenotypes we observed. We used the *Efcab9-28del* mutant line as the *Efcab*9-null mouse throughout this study.

**Figure 2.**
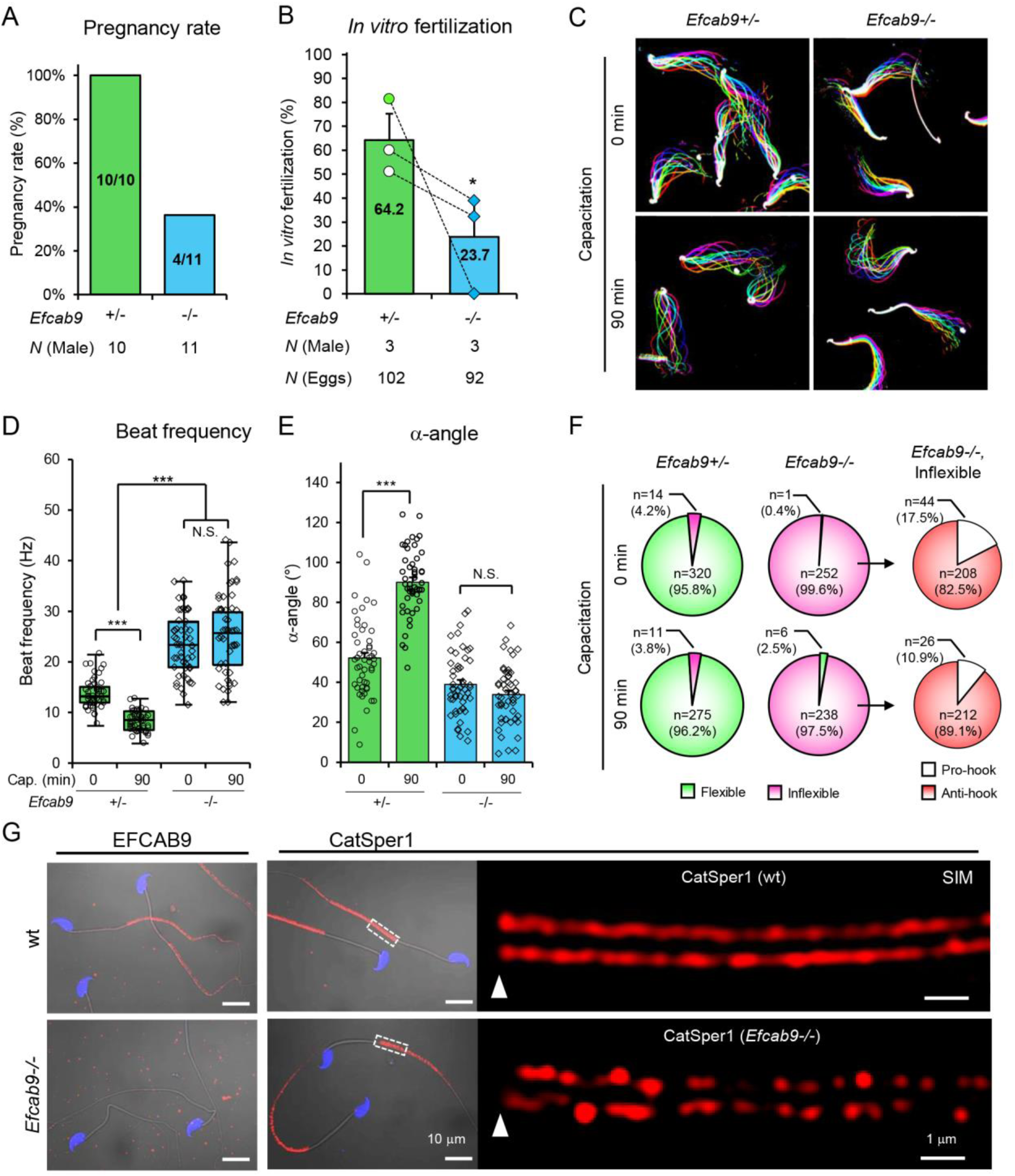
Genetic disruption of *Efcab9* compromise male fertility, sperm motility, and CatSper domain. (A) Percent pregnancy rate of *Efcab9*-/- compared with *Efcab9*+/- males over two months when housed together with wt females. (B) *In vitro* fertilization rate of sperm cells from *Efcab9*+/- (64.2 ± 11.0%) and *Efcab9*-/- (23.8 ± 14.7%) males with cumulus oocyte complex. \**P*<0.05. (C) Flagellar wavefrom of *Efcab9*+/- (left) and *Efcab9*-/- (right) sperm cells. Movies recorded at 200 fps from the cells tethered to glass coverslips before (top, 0 minute) and after (bottom, 90minutes) incubating under capacitation conditions. Overlays of flagellar traces from two beat cycles are generated by hyperstacking binary images; time coded in color. (D) Beat frequency before (0 minute, *Efcab9*+/-, 13.9 ± 0.4 Hz; *EFCAB9*-/-, 23.5 ± 0.9 H z) and after (90 minutes, *Efcab9*+/-, 8.19 ± 0.27 Hz; *Efcab9*-/-, 26.2 ± 1.2 H z) incubating under capacitation conditions is shown box plot, indicating quartile distribution. Median ± SEM, N=50 each group. \*\*\**p* <0.001. (E) The maximal bending angle of midpiece in primary anti-hook curvature was measured from *Efcab9*+/- (0 minute, 52.1 ± 2.7°; 90 minutes, 90.1 ± 2.3°) and *Efcab9*-/- (0 minute, 40.0 ± 2.2; 90 minutes, 33.8 ± 2.1°) sperm, and tail parallel to head is considered to 0°. Data is represented as mean ± SEM. N=50 for each group. \*\*\**p* <0.001 (F) Flexibility and flagellar bending patterns of sperm cells from *Efcab9*+/- (left) and *Efcab9*-/- (middle and right) mice. The flexibility of sperm tails is examined and classified in to flexible (green) or inflexible (magenta). Inflexible tails are further divided into pro-hook (white) or anti-hook (red) according to the direction of the flagellar bend. Sperm numbers examined from three independent experiments are shown in the pie charts. (G) *Efcab9*-/- spermatozoa have normal sperm morphology but display discontinuous CatSper domain. Overlay of confocal images and the corresponding DIC images of cells of α-EFCAB9 (left) and α-CatSper1 (right) immunostained sperm cells from wt (upper) and *Efcab9*-/- (lower) mice. The inset areas were subject to SIM microscopy as shown at the right side of the CatSper1 confocal images. *See also Figures S3 and S4; Video S1*.

The resulting frame-shifted transcripts failed to generate EFCAB9 proteins, demonstrated by immunoblotting and immunocytochemistry (Figures S3C and SG). Mutant mice lacking EFCAB9 display no gross phenotypic abnormality in appearance, behavior, or survival. *Efcab9-null* females had normal mating behavior and gave birth to litters when mated with wt or mono-allelic heterozygous (het) males. However, fertility was severely impaired in Efcab9-null male mice (Figures 2A, 2B, and S4A), despite normal sperm morphology and epididymal sperm count (Figures 2G and S4B). The pregnancy rate of the null males was 36% when housed together with fertile females over 2 months (Figure 2A), compared to 100% for wt. The fertilization rate of Efcab9-null spermatozoa was also significantly reduced *in vitro* (Figure 2B). Finally, *Efcab9*-null males that sire pups tend to have smaller average litter size than Efcab9-het males (Figure S4A), suggesting that Efcab9-null males are subfertile.

These results prompted us to further examine the flagellar beating pattern of head-tethered spermatozoa (Figure 2C; Video S1). Incubating under capacitating conditions increased the amplitude of lateral movement, measured as the maximum angle within the midpiece (α-angle) (Qi et al., 2007) and slowed the beat frequency of *Efcab9*-het sperm (Figures 2D and 2E; Video S1). In contrast, the beat frequency of Efcab9-null sperm cells, which is faster than *Efcab9*-het spermatozoa, did not respond to capacitating conditions. Importantly, we observed that *Efcab9*-null spermatozoa displayed rigid proximal flagella with a fixed midpiece curvature (Figures 2C and 2F; Video S1). The majority of the Efcab9-null sperm remained bent in the anti-hook direction (Ishijima et al.,2002) (Figures 2C and 2F), suggesting that EFCAB9 functions in the CatSper-mediated Ca^2+^ signaling pathway that normally dominates and results in the pro-hook bend (Chang and Suarez, 2011). Supporting this idea, capacitation-associated protein tyrosine phosphorylation was elevated in *Efcab9*-null spermatozoa (Figure S4C), indicating that CatSper-mediated Ca^2+^ current was aberrant as previously demonstrated by genetic disruption (Chung et al., 2017; Chung et al., 2014) or pharmacological inhibition (Navarrete et al., 2015).

These findings led us to investigate whether EFCAB9 deficiency dysregulates the localization and/or expression of the CatSper channel complex in sperm cells. SIM images of CatSper1 revealed that the structural continuity of the linear arrangement of CatSper channels is fragmented in the absence of EFCAB9 while the CatSper complex remains targeted to the flagellum (Figure 2G). We examined the protein expression of CatSper subunits in *Efcab9*-null sperm and found that all the transmembrane (TM) CatSper subunits express at 20-30% of normal (Figure S3). All these results are identical to the previously reported *CatSperz*-null phenotypes (Chung et al.,2017). Thus, we set out to determine if EFCAB9 and CatSperζ form a functionally-relevant binary complex.

### EFCAB9 Forms a Binary Complex by Interacting Directly with CATSPERζ

Comparing Efcab9-null mice with the previously established *CatSper1-,d-*, and *z*-knockout lines, we discovered that EFCAB9 and CatSperζ protein expression is strictly interdependent (Figures 3A and 3B). In contrast, neither is required for CatSper channel expression (Figures 2G, 3A, and 3B), but each modulates the protein expression levels and organization of CatSper Ca^2+^ signaling domains (Figure 2G) (Chung et al., 2017). To further assess their functional interaction, we generated double knockout males (*Efcab*9-/-; *CatSperz*-/-). We did not observe any difference in defects of double knockout males (Figures D-S4F) compared with those of single *Efcab9*- or *CtSperz*-null males (Figure 2) (Chung et al., 2017). These mice were severely subfertile with ~33% pregnancy rates in mating studies (Figure S4E). Sperm from double knockout male mice failed to develop hyperactivated motility after incubation under capacitating conditions. Their sperm display flagellar waveforms similar to those observed in sperm from single *Efcab9*- or *CatSperz*-null male mice in that double knockout sperm also show the inflexible proximal tail and reduced flagellar waveform amplitude (Figures 2C-2F and S4F) (Chung et al., 2017). Sperm from double knockout males express CatSper proteins comparable to those of Efcab9-null sperm cells, showing thinning of the linear CatSper domains and loss of continuity (Figures S4G and S4H). These results demonstrate no additive or synergistic defects in the loss of both proteins in male germ cells, strengthening the conclusion that EFCAB9 and CatSperζ are dispensable for CatSper channel expression.

**Figure 3.**
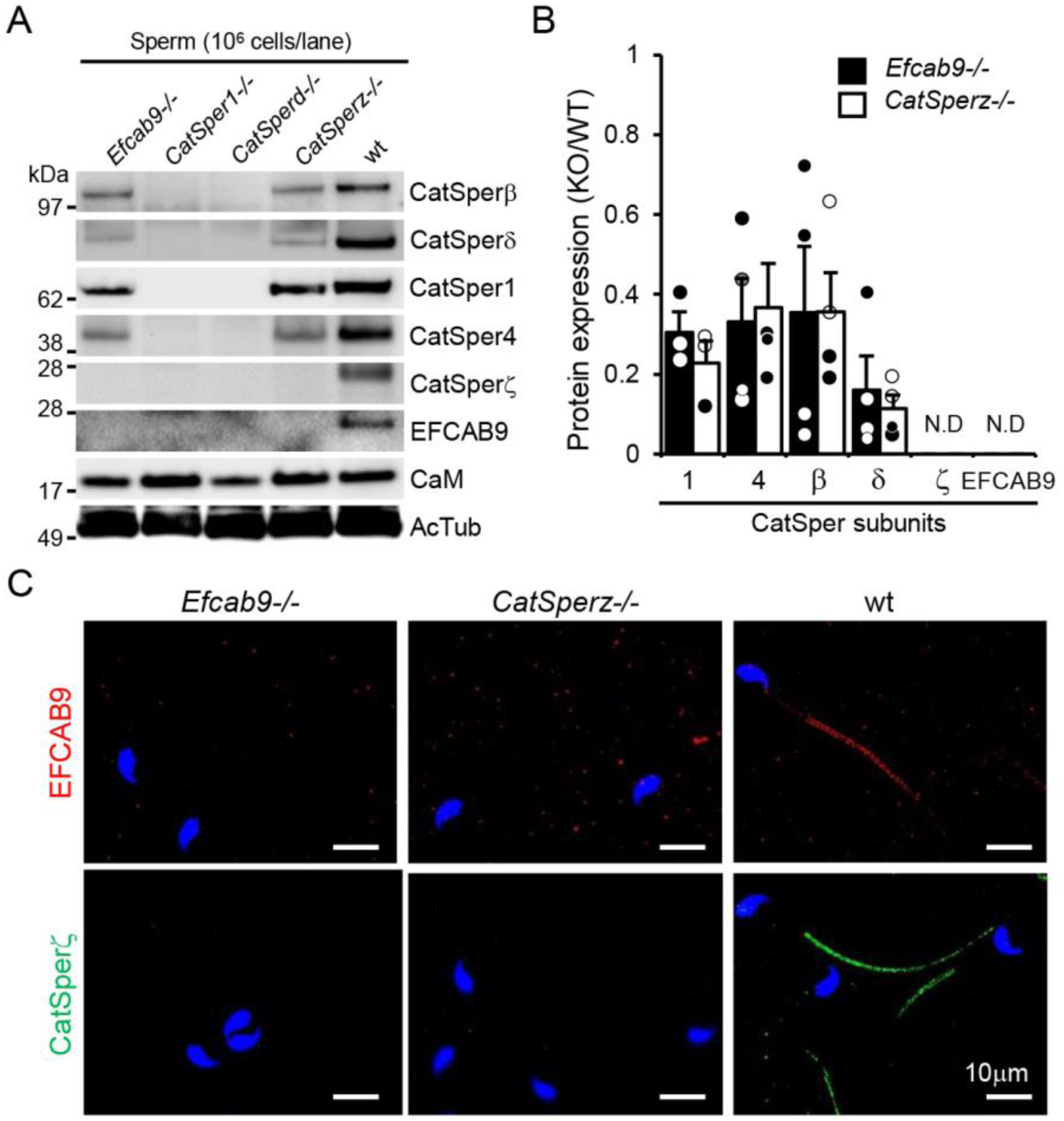
EFCAB9 and CatSperζ form a binary complex in spermatozoa. (A-B) EFCAB9 and CatSperCatSperζ proteins express interdependently in spermatozoa. (A) T he absence of either EFCAB9 or CatSperζ is dispensable for the protein expression of all the other CatSper subunits with transmembrane domains but eliminates the expression of each other. (B) Protein expression levels of CatSper subunits in *Efcab9*-/- and *CatSperz*-/- sperm cells compared with those of wt sperm. The relative levels of CatSper subunits in *Efcab9*-/- vs. *CatSperz*-/- sperm are all comparable; CatSper1 (0.31 ± 0.05 vs 0.23 ± 0.06), CatSper4 (0.33 ± 0.11 vs 0.37 ± 0.11), CatSperβ (0.35 ± 0.16 vs 0.36 ± 0.10), CatSperδ (0.16 ± 0.08 vs 0.11 ± 0.03), CatSperζ and EFCAB9 (not detected). Data is represented as mean ± SEM, N=3. (C) Confocal images of immuno sta ined EFCAB9 (red) and CatSperζ (green) in *Efcab9*-/-, *CatSperz*-/-, and wt spermatozoa. Hoechst dye stains sperm head (blue). See *also Fig ure S4*.

To determine whether EFCAB9 binds CatSperζ, we carried out immunoprecipitation and pull-down analysis. For immunoprecipitation experiments, 293T cells were transiently transfected with plasmid(s) encoding either EFCAB9 or CatSperζ alone, or both EFCAB9 and CatSperζ. As seen in Figure 4A, EFCAB9 and CatSperζ both co-immunoprecipitated (Co-IP) the other protein. To test for a direct interaction between EFCAB9 and CatSperζ, recombinant mouse EFCAB9 and CatSperζ proteins fused to either Glutathione S-transferase (GST) or 6xHis-GB1 were purified from *E. coli* and used for *in vitro* binding assays. GST or His-tag pull-down analysis revealed a strong interaction between EFCAB9 and CatSperζ(Figure 4B), suggesting that EFCAB9 and CatSperζ are direct binding partners.

**Figure 4.**
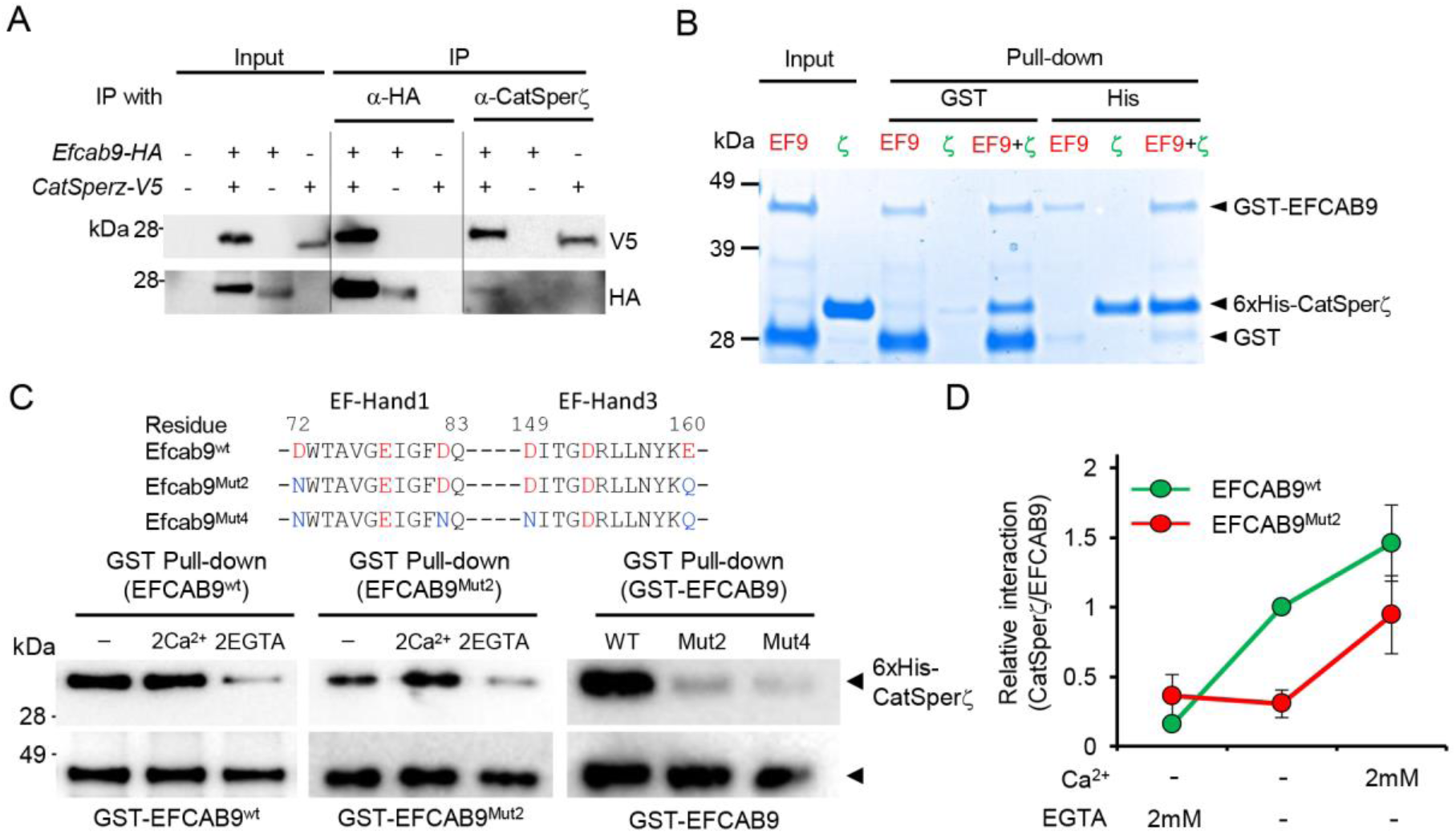
EFCAB9 directly binds to CatSperζ and the interaction is Ca^2+^-sensitive. (A) Co-immunoprecipitation of EFCAB9 with Catseprζ. HEK293T cells transfected with plasmids encoding HA-tagged EFCAB9 and/or V5-tagged CatSperζ were immunoprecipitated with anti-HA or anti-CatSperζ and the immunocomplexes were blotted with anti-HA or V5 antibodies. (B) Pulldown analysis of purified recombinant EFCAB9 and CatSperζ proteins. GST or His pull-down analysis was performed between full-length GST-EFCAB9 and 6xHis-tagged CatSperζ fused to Gb1. EFCAB9 and CatSperζ interact directly. (C) Ca^2+^ dependent interaction between EFCAB9 and CatSperζ through the EF hands. The mutants used are indicated as EFCAB9^Mut2^ (D72N/E1 60Q) and EFCAB9^Mut4^ (D72N/D82N and D149N/E160Q), respectively. GST pull-down analysis was performed between GST-EFCAB9 (left) or GST-EFCAB9^Mut2^ (middle) and 6xHis-CatSperζ in medium with nominal free Ca^2+^ (-), 2 mM CaCl_21_ (2Ca^2+^), or nominally calcium-free plus 2 mM EGTA (2EGTA) at pH 7.5. The effect of mutations in the EF hands of EFCAB9 on the interaction with 6xHis-CatSperζ (right) was analyzed by GST pull-down assay in the medium with nominal free Ca^2+^ at pH 7.5. Data is represented as mean ± SEM, N=5. (D) Relative amount of 6xHis-CatSperζ directly bound to GST pull-down EFCAB9 in various free calcium concentrations in (C). Binding buffer with nominal free Ca^2+^ is without adding any additional calciumions and contains ~ 10μM free Ca^2+^ as a trace ion. All the recombinant proteins in (B, C, and D) were purified from *E. coli*.

### The EF-hands of EFCAB9M ediate Ca^2+^-sensitive Interaction with CATSPERζ

Next, we examined whether EFCAB9-CatSperζ interaction is Ca^2+^-sensitive to test the predicted Ca^2+^ binding ability of EFCAB9. GST pull-down analysis revealed a strong interaction between EFCAB9 and CatSperζ under conditions without added Ca^2+^ (nominal free Ca^2+^, typically ~ 10 μM free Ca^2+^) or with 2 mM Ca^2+^ at pH 7.5 (Figures 4C, left, and 4D). In contrast, the EFCAB9 and CatSperζ dissociate, and binding is minimal when Ca^2+^ is chelated by addition of 2 mM EGTA. These results indicate that Ca^2+^ is required for maintaining a stable EFCAB9-CatSperζ interaction and that Ca^2+^ likely binds to the EF-hand domains of EFCAB9.

To test whether the Ca^2+^ sensitive interaction is due to the Ca^2+^ binding ability of EFCAB9, we generated two EFCAB9 mutants by changing conserved, negatively-charged residues in the EF-hand motifs (Figures 1B and 4C). We substituted the conserved aspartate at position 72 in EF1 to asparagine and the glutamate at position 160 in EF3 to glutamine (D72N/E160Q, EFCAB9^Mut2^) by site-directed mutagenesis.EFCAB9^Mut4^ was produced by additional mutations of aspartate at positions 82 in EF1 and 149 in EF3 to asparagine (D72N/D82N and D149N/E1 60Q) (Figures 1B and 4C). These two charge-neutralizing mutants were then used for *in vitro* binding assays. Diminished EFCAB9^WT^ binding to CatSperζ by chelating free Ca^2+^ (Figure 4C, left) suggests that EFCAB9 interacts with CatSperζ in its calcium-bound form. Since EFCAB9^Mut2^ has significantly reduced binding to CatSperζ (Figures 4C and 4D), D72N/E160Q mutations located at the end s of the EF1 and EF3 loops attenuate Ca^2+^ binding of EFCAB9. Ca^2+^ binding to EFCAB9^Mut2^ is partly rescued by adding 2 mM Ca^2+^ (Figure 4C; middle, and 4D). Substitution of the four acidic residues to neutralizing amino acids (D72N/D82N and D149N/E160Q, EFCAB9^Mut4^), however, almost eliminated EFCAB9 binding to CatSperζ (Figure 4C, right), suggesting that EF2 has a lesser contribution to Ca^2+^ binding. These results confirm that the EF-hands of EFCAB9 mediate Ca^2+^ sensitive interaction, supporting the conclusion that EFCAB9, in complex with CatSperζ, can function as a Ca^2+^ sensor for the CatSper channel. Thus EFCAB9-CatSperζ appears to be im porta nt for both modulating channel activity and organizing the CatSper domains.

### EFCAB9-CatSperζ Is Required to Link Two Rows within a Single CatSper Domain

CatSper channels form a highly organized Ca^2+^ distribution network in four quadrants along the principal piece of sperm flagella, which control and coordinate Ca^2+^ signaling along the extremely long, narrow tail during hyperactivated motility (Chung et al., 2017; Chung et al., 2014. Our previous silver-intensified immunogold electron microscope (EM) images of CatSper 1 Chung et al., 2014) suggested a two-row structure within each longitudinal CatSper compartment. However, a typical transmission EM shows only a subsection of a flagellum and standard STORM imaging with 20- to 50-nm 3D resolution did not clearly resolve the paired structure. To further assess the potential substructure and the effect of EFCAB9-CatSperζ on thinning at repeated intervals in the absence of EFCAB9(Figure 2G) and/or CatSperζ (Figure S4H) (Chung et al., 2017 we employed the recently developed, 4Pi single-molecule switching nanoscope (4Pi-SMSN) that enables improved resolution of 10-to 20-nm in three dimensions (Huang et al., 2016).

We first imaged CatSper domains in wt spermatozoa by immunolabeling with a verified CatSper1 antibody (Chung et al., 2014; Ren et al., 2001). Consistent with our previous findings, we visualized the CatSper domains as distinct linear quadrants along the principal piece of the flagella (Figures 5A and 5B; Video). Closer inspection by 4Pi-SMSN revealed that each CatSper domain is further resolved to two rows, as clearly demonstrated by the cross-section image and the angular distributions (Figure A and 5B; Video). Interestingly, the CatSper domains are not only disrupted at periodic intervals, but the two-row structure within a domain appears to be more irregular in the absence of EFCAB9-CatSperζ(Figures 5A and 5B; Video S3). These results illustrate that CatSper double-row organization requires EFCAB9-CatSperζ.

Micron-scale membrane subdomains have been demonstrated in terms of lipid segregation in the sperm head (Selvaraj et al., 2006). In sperm tails, a distinct membrane subdomain known as the flagellar zipper was reported (Friend and Fawcett, 1974; Selvaraj et al., 2007) but the function and molecular basis of this zipper structure remains a mystery. We asked whether sperm tail membrane subdomains, representing the macromolecular CatSper complex, can be visible by scanning EM (SEM). We identified a doublet of raised linear surface domains in wt mouse sperm, running each side of the longitudinal columnar surface structure down the principal piece (Figure 5C, top). One doublet is made of two ∽20 nm thick membranous stripes. The absence of these raised stripes in the SEM images of *CatSper1*-null spermatozoa (Figure 5C, bottom) indicates that these linear surface domains can be a protein-based membrane compartmentalization composed of the CatSper channel complex. The shorter fragmented singlet observed in *Efcab9*-null sperm flagella (Figure 5C, middle) supports the conclusion that EFCAB9-CatSperζ has a structural function in organizing doublet CatSper domains seen in the sperm surface nanoarchitecture.

**Figure 5.**
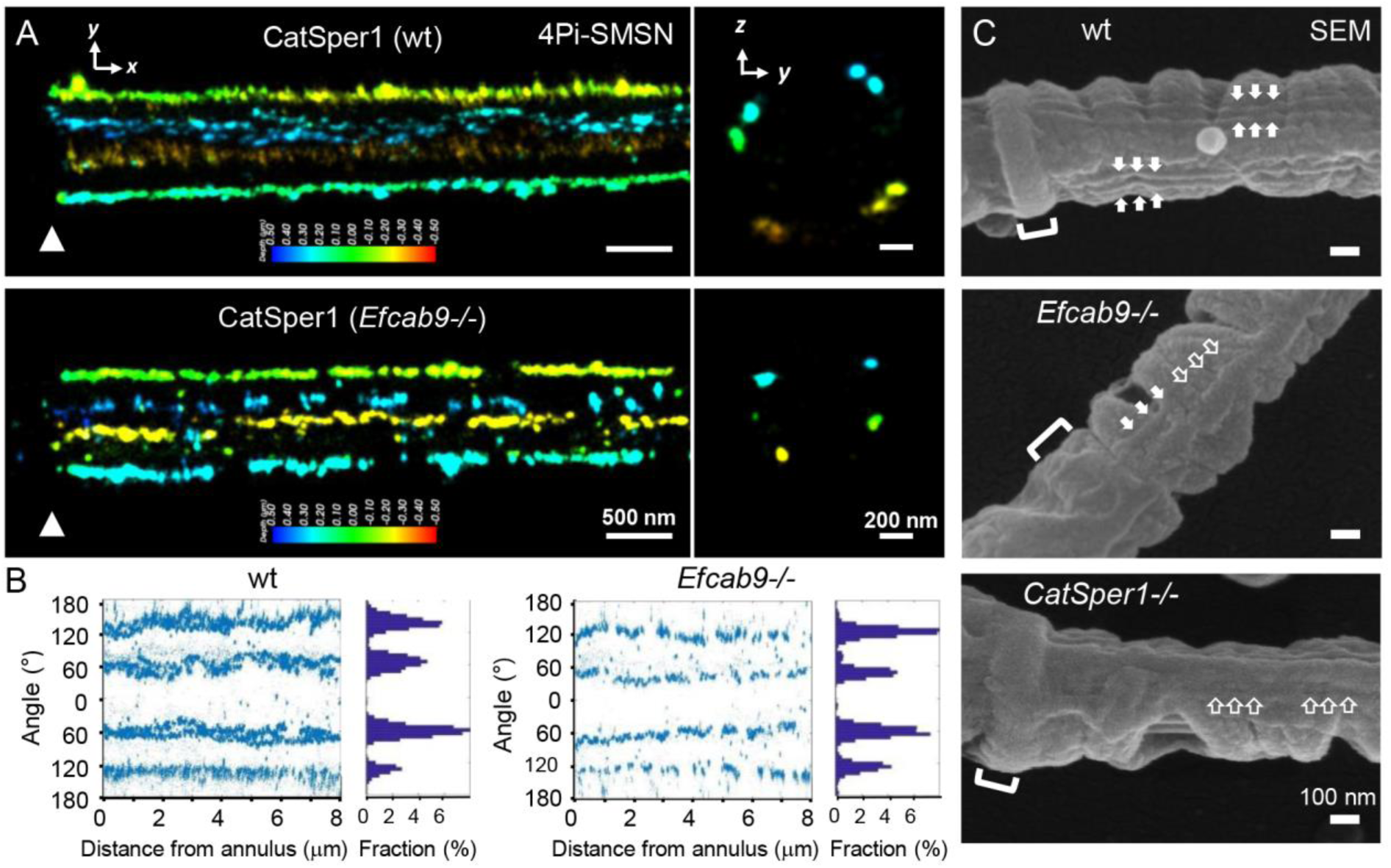
Loss of EFCAB9 alters the two-row structures of CatSper domains and sperm surface nanoarchitecture. (A) 3D 4Pi-SMSN images of CatSper1 in wt (top) and *Efcab9*-/- (bottom) flagella. *x-y* projection colors encode the relative distance from the focal plane along the *z* axis. Scale bar, 500 nm. Arrowheads in each panel indicate annulus. *y-z* cross sections (100 nm thick) at ~ 3 um from the annulus are shown on the right. Scale bar, 200 nm. (B) Angular distributions (left) and profile (right) of the surface-localized molecules of 4Pi-SMSN images of CatSper1 in wt (upper) and *Efcab9*-/- (lower) shown in (A). Two-rows are seen in each CatSper quadrant in wt but only single line in *Efcab9-/-* spermatozoa. (C) Scanning electron microscopic images sperm flagella from wt (top), *Efcab9*-/- (middle), and *CatSper1*-/- (bottom) mice. Two-row structure organization on the membrane surface are observed in both sides of longitudinal column in wt sperm but only one fragmented line in *Efcab9*-/- spermatozoa. No linear presentation is observed in *CatSper1*-/- sperm cells. Scale bar, 100 nm. *See also Video S2 and S3*.

### EFCAB9-CatSperζ Complex Confers pH-Dependent Activation and Ca^2+^ Sensitivity to CatSper Channel

A rise in intracellular pH potentiates *I_CatSper_* (Kirichok et al., 2006 which regulates sperm motility. In light of the reduced average whole cell *I_CatSper_* amplitude in *CatSperz*-null spermatozoa (Chung et al., 2017) together with the new findings of the Ca^2+^ dependent binary complex formation of EFCAB9-CatSperζ (Figures 3, 4 and S4 we next examined whether the loss of EFCAB9 and CatSperζ altogether affects the pH dependent activation and cytosolic Ca^2+^ sensitivity of the CatSper current upon voltage changes. A reduced current can reflect changes in the number, open probability (*P_o_*), or elementary conductance of the channels. Given peculiarities of reduced protein expression of other companion CatSper subunits in *CatSperz*-null, *Efcab9*-null, and double knockout spermatozoa, we characterized *I_CatSper_* densities (pA/pF) (Figures 6A-6F and S5).

Interestingly, with genetic disruption of *CatSperz,* which results in spermatozoa lacking both EFCAB9 and CatSperζ, current densities are similar to wt (pH 6.0) (Figure S5A-S5D). At higher negative and positive voltages, CatSper currents are even slightly larger, but since these are outside the physiological range, are of uncertain significance. In agreement with previous findings (Kirichok et al., 2006), *I_CatSper_* densities increase dramatically in response to addition of 10 mM NH_4_Cl in wt spermatozoa (Figure S5). In contrast, the response to addition of 10 mM NH_4_Cl was only modest in *CatSperz*-null sperm cells (Figure S5 suggesting a compromised channel activation by intracellular alkalinization. These results provide evidence that EFCAB9-CatSperζ regulates pH-dependent CatSper activation.

Low free Ca^2+^ (<10 nM, buffered by 2 mM EGTA) largely eliminated the interaction between purified recombinant EFCAB9 and CatSperζ *in vitro* (Figures 4C and 4D). To directly test the Ca^2+^ sensitivity of the pH-dependent CatSper activation, we recorded the currents with buffered [free-Ca^2+^] at a fixed intracellular pH in pipette (Figures 6A-6F). At acidic intracellular pH (pH_i_, = 6.0), CatSper channel was mainly closed in the inward direction in both wt and Efcab9-null sperm during voltage step regardless of intracellular Ca^2+^ concentrations. This result suggests that the apparent Ca^2+^ sensitivity of the CatSper channel is low at acidic pH, preventing channel full activation. However, alkaline intracellular pH (pH_i_, = 7.4) dramatically potentiated CatSper channel activation in wt spermatozoa (20-fold) but to a much less degree in *Efcab9*-null sperm (~ 5 fold) particularly in the presence of 10 μM intracellular free Ca^2+^ (Figures 6A-6F), which corresponds to calcium levels in the vicinity of calcium channels upon Ca^2+^ entry (Naraghi and Neher, 1997). Thus, pH presumably tunes the Ca^2+^ sensitivity of EFCAB9, triggering a conformational change, which change the affinity of EFCAB9 for CatSperζ within the binary complex.

To better understand the pH-dependent Ca^2+^ gating mechanisms of CatSper by EFCAB9-CatSperζ, we performed GST pull-down analysis of EFCAB9/CatSperζ interactions at varying pH using purified recombinant proteins in nominal free calcium solution without adding any additional calcium ions (Figures 6G and 6H). Usually, such solutions contain about 10 μM free calcium as a trace ion. We found that the amount of CatSperζ bound to wt EFCAB9 gradually decreases when pH is raised (Figures 6G, left, and 6H). T he interaction is further diminished when GST pull-down was performed with EFCAB9^Mut2^ (Figure 6G, right, and 6H). These data support our conclusion that the EFCAB9 is a pH-dependent Ca^2+^ sensor to activate CatSper channel, indicating that evolution developed ways to limit CatSper-mediated Ca^2+^ entry before capacitation-associated intracellular alkalinization in mammals.

**Figure 6.**
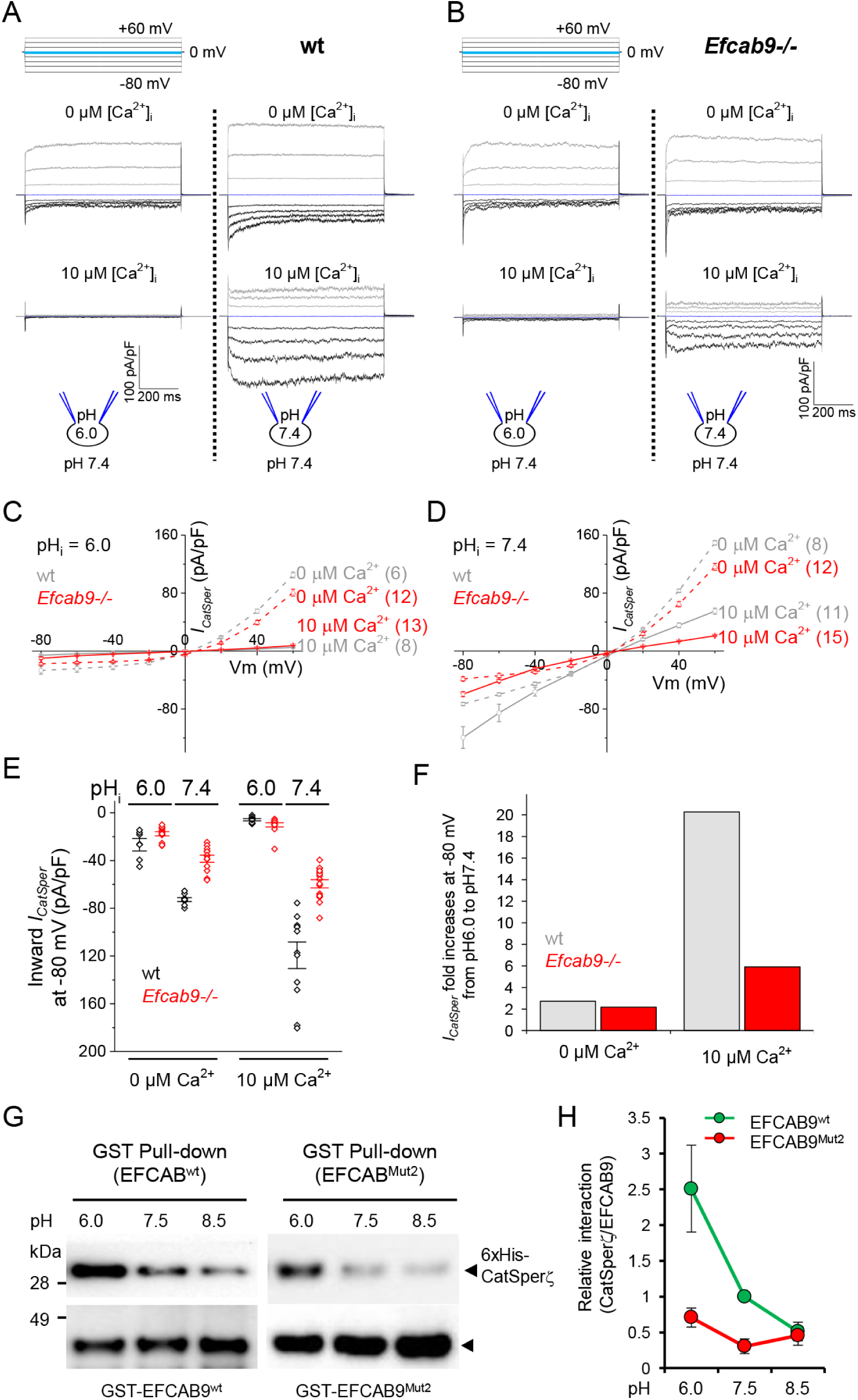
EFCAB9 confers maximal CatSper channel activity in a pH- and Ca^2+^-dependent manner, (A-B) Representative *I_CatSper_* traces from wt (A) and *Efcab9*-/- (B) spermatozoa recorded with 0 μM (top) or 10 μM (bottom) free intracellular calcium at intracellular pH = 6.0 (left) or intracellular pH = 7.4 (right). Currents were elicited by a step protocol from- 80 mV to +60 mV in 20 mV increments and a holding potential of 0 mV as shown. The cartoon indicates the intracellular and bath pH used for each panel. (C-D) I-V relationship of *I_CatSper_* traces deducted at steady state levels from recordings of wt (gray lines) and *Efcab9*-/-(red lines) spermatozoa with 0 pM (dashed lines) or 10 μM (solid lines) free intracellular calcium at pH_i_ = 6.0 (C) and pH_i_ = 7.4 (D). (E) Averaged inward *I_catsper_* from wt (black) and *Efcab9*-null (red) sperm collected at −80 mV at respective experimental conditions. (F) Bars represent fold increase of the inward *I_CatSper_* at-80 mV when switching from intracellular pH 6.0 to pH 7.4 in wt (grey) and Efcab9-null (red) spermatozoa. (G) EFCAB9-CatSperζ complex dissociates when pH increases from 6.0 to 8.5. GST pull-down analysis was performed between GST-fused EFCAB9 (left) or EFCAB9^Mut2^ (right) and 6xHis-tagged CatSperζ under the increasing pH conditions at pH 6.0, 7.5, and 8.5. Binding buffer contains ~ 10 pM free Ca^2+^ as a trace ion (nominal free Ca^2+^ without adding any additional calcium ions). (H) Quantitation of relative CatSperζ bound to EFCAB9. N=5. All the recombinant proteins were purified from *E. coli* as in *Figure 4*. Data is represented as mean ± SEM (C, D, E, and H); (n) indicates the number of individual cells used. See *also FFigure S5*.

### EFCAB9-CatSperζ Interacts with Cytoplasmic Mouth of CatSper Channel Pore

Presumably due to the complex composition and the organization into highly ordered arrays (Figures 3 and 5), native CatSper channel complex is tightly associated with the cytoskeletal structures and not solubilized from mature sperm cells (*unpublished* data). Therefore, we examined molecular interaction between EFCAB9-CatSperζ and individual CatSper subunits by performing Co-IP in a heterologous system and pull-down assays using purified recombinant proteins (Figures 7A and 7B and S6A-F). HA-tagged EFCAB9 and V5-tagged CatSperζ encoded by a bi-cistronic plasmid were transiently expressed in 293T cells with either one of the CatSper subunits. Co-IP revealed that the EFCAB9-CatSperζ complex interacts with each of CatSper1-4 (Figures 7A and S6A-S6C) but has little to no interaction with the auxiliary transmembrane subunits (β, γ δ, and ε) (Figures 7B), suggesting that EFCAB9-CatSperζ interacts mainly with the channel pore. The remarkably histidine-rich N-terminus of CatSper1 was suggested as a candidate for the pH-sensor of CatSper channel (Kirichok et al., 2006; Ren et al., 2001 yet our electrophysiology results strongly suggest that the EFCAB9-CatSperζ complex can account for the pH-dependence of channel activation (Figures 6 and S5). Interestingly, pairwise-distance analysis of amino acid sequences showed that the CatSper1 N-terminal domain and CatSperζ are highly divergent among mammalian species in contrast to EFCAB9, the C-terminal domain of CatSper1 and the cytoplasmic domains of other CatSperα subunits (Figures 7C and S6G and S6H), suggesting specific co-evolution. To further examine the potential domain interaction between CatSper1 and EFCAB9-CatSperζ, we performed pull-down analysis between recombinant amino (N1-150) or the carboxyl (C574-686) terminus of CatSper1 and the recombinant CatSperζ or EFCAB9-CatSperζ complex (Figures S6D-S6F). We found no evidence of direct interaction between either the carboxyl or amino terminal domain of CatSper1 and EFCAB9-CatSperζ or CatSperζ (Figures S6E and S6F). This does not, however, rule out the possibility that the N-terminus has a separate pH-dependent effect on gating, or that it binds Zn^2+^ or other modulators.

**Figure 7.**
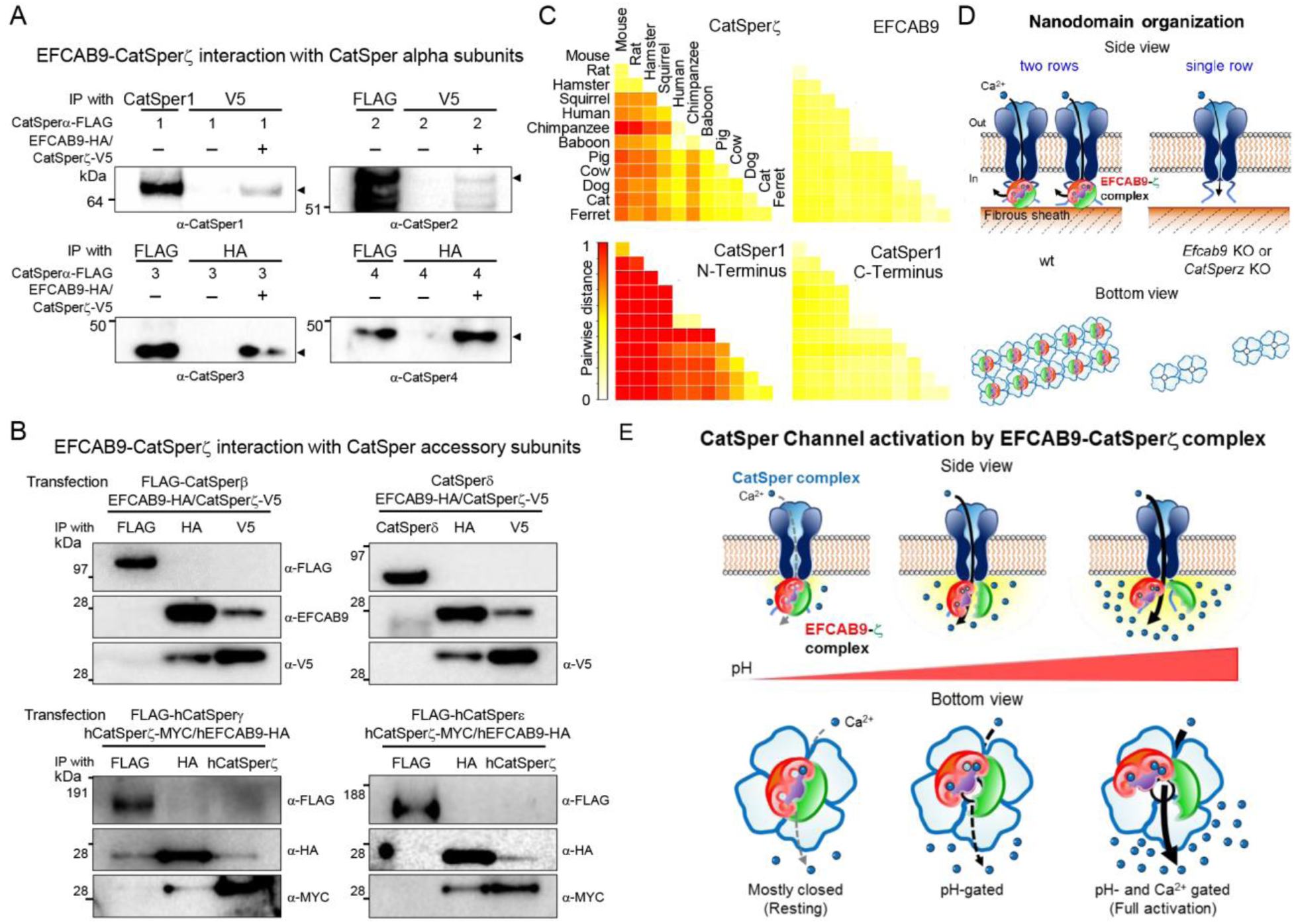
Molecular interaction of EFCAB9-CatSperζ binary complex with other CatSper subunits. (A) Co-immunoprecipitation assays were performed from 293T cells transfected with plasmids encoding CatSperζ-V5-(P2A)-EFCAB9-HA and one of each CatSper alpha subunit (1, 2, 3, or 4). After immunoprecipitation with anti-HA or anti-V5 resin, precipitates were analyzed by immunoblotting with antibodies against each CatSper protein to detect the interaction to EFCAB9-CatSperζ complex. (B) Co-immunoprecipitation assays were performed with each accessory (β, γ, δ, or ε) subunit with EFCAB9-CatSperz complex. Reciprocal interaction was analyzed by antibodies against each protein. (C) Conservation of CatSperζ, EFCAB9, and N-and C-terminal domains of CatSper1 amino acid sequences among mammals based on a pairwise-distance analysis. The color scheme in heat maps represents the levels of sequence variation scored by pairwise distance between two orthologues. The value of pairwise distance over 1 was set to 1 (red). (D) CatSper nanodomain organization in wt and EFCAB9 and/or CatSperζ deficient sperm. (E) Model for CatSper channel activation by EFCAB9-CatSperζ binary complex. pH_i_ elevation during capacitation partially dissociates EFCAB9 from CatSperζ, which releases gate inhibition and opens the pore, enabling EFCAB9 to bind entering Ca^2+^ and undergo a conformational change to maintain the prolonged open state of the channel.See *also FFigure S6*.

## Discussion

We propose a molecular mechanism regulating domain organization and pH-dependent activation of the CatSper channel (Figures 7D and 7E). Functional characterization of EFCAB9 in complex with CatSperC links the CatSper downstream Ca^2+^ signaling pathway and sperm hyperactivated motility. Our results show that EFCAB9, a protein missing in the *CatSper1*-null sperm lacking CatSper channel expression, is an evolutionarily conserved CatSper channel component. EFCAB9 is not required for functional CatSper channel formation. Instead, EFCAB9 forms a binary complex with CatSperC through a direct and Ca^2+^ sensitive interaction. The EFCAB9-CatSperζ complex organizes the doublet CatSperζ a^2+^ signaling domains. Mutant CatSper channels lacking EFCAB9-CatSperζ largely failed to respond to a rise in intracellular pH, leading to severe male subfertility. One possibility is that calcium entering via CatSper rapidly binds EFCAB9 that, by altering its association with the channel complex, maximizes flagellar shape and motility change in adaptation to hyperactivating conditions. In non-hyperactivating conditions, such as when pH is low, CatSperζ-complexed EFCAB9 limits calcium entry via CatSper.

### Significance of the CatSper Proteome Screen and Identification of EFCAB9

The identification of all the nine known CatSper subunits and EFCAB9 in the strong downregulation cluster’ suggests that the screen is presumably near saturation. The lack of significant change in overall protein expression in *CatSper1*-null compared to wt spermatozoa suggests that there is no major compensatory protein expression induced by the loss of the CatSper channel complex.

Our comparative screen identified many EF-hand containing proteins, which include Centrin1, EFCAB1, 2, 3, 5, 6, 9, 10, EFHB, EFHC2, PLCζ, PPEF1, LETM1, and CaM (Table S1). Among these, only EFCAB9 showed significant change with the loss of CatSper subunits. Since the EF-hand is one of the major Ca^2+^-binding regions found on many of the Ca^2+^ sensors, we propose that EFCAB9 is specialized for the CatSper channel. Intriguingly, the *Efcab9* gene shares the unique pattern of lineage-specific gains and losses of other CatSper genes (CatSper1-4, CatSperβ, γ, and ε) from single-celled eukaryotic flagellates (FFigure S1). This suggests that *Efcab9* is a part of the set of core genes comprising an evolutionarily conserved unit of flagellar Ca^2+^ channel.

The EF hand calcium binding motif contains canonical 12-residues crucial to coordinate the calcium ion (Gifford et al., 2007). EF-hands tend to occur in pairs, which form a discrete domain. For example, the classical sensor CaM has four EF-hands that comprise N- and C-terminal lobes; each lobe contains a pair of EF-hands (N-lobe: CaM-EF1/2 and C-lobe. CaM-EF3/4, FFigure S2A). Ca^2+^ binding induces a con fo rmational change in the EF-hand motif, influencing interaction with target proteins. EFCAB9 is a rare example of odd-numbered EF-hand proteins (Figure 1B). The EF-hand corresponding to CaM’s EF1 is absent, resulting in a single, non-canonical EF-hand in the N-lobe. Here, glutamate is substituted by glutamine or lysine at the twelfth residue in various species including mouse and human (EFCAB9-EF1, Figures 1B and S2C). Substitution at this position was previously shown to disable Ca^2+^ binding in the CaM EF-hand motif (Beckingham, 1991). In mammals, the first EF-hand in the EFCBA9 C-lobe also lacks glutamate at the EF loop’s twelfth position (EFCAB9-EF2). We surmise that this enables EFCAB9 to bind CatSperζ, a mammalian-specific CatSper component, albeit with a loss in Ca^2+^ affinity.

### EFCAB9 is a Ca^2+^ Sensor for CatSper Channel

Cytoplasmic calcium binds directly to channel proteins, or through adaptor or modulatory proteins, to regulate their activity (Clapham, 2007; Yu and catteral, 2004). Examples of the intrinsic Ca^2+^ sensors include the voltage-gated Ca^2+^ channels, Ca_v_1.2 (Van Petegem et al., 2005) and the *Arabidopsis thaliana* two pore channel, TPC1 (Guo et al., 2016), as well as RCK-motifs in calcium-regulating BK potassium channels ((Piskorowski and Aldrich, 2002). EF-hands are found within the cytoplasmic regions of these channel proteins and alter channel function upon localized calcium accumulation. In contrast, the small conductance Ca^2+^ activated K+ channel, SK (Lee and MacKinnon, 2018) and the subfamily of KCNQ voltage-gated K^+^ channels, Kv7.4 (Chang et al., 2018) forms a complex with Ca^2+^ binding protein, CaM, which regulates Ca^2+^ dependent activation.

None of the cytoplasmic regions of the nine CatSper subunits contains known calcium/CaM binding motifs. In the current study, we report EFCAB9 as a Ca^2+^ sensing auxiliary subunit for CatSper channel and establish a regulatory mechanism based on the interaction with the binary EFCAB9-CatSperζ complex. Our combinatorial mutagenesis studies reveal that EF1 and/or EF3 of EFCAB9 bind Ca^2+^. The contribution of individual EF-hands in Ca^2+^-binding and/or the interaction with CatSperζ will require further investigation.

In the absence of active CatSper-mediated Ca^2+^ entry before capacitation, EFCAB9-CatSperζ pre-associates with the channel at basal levels of intracellular (~100 nM free) calcium. In this resting state, the channel pore would be mostly closed. In the presence of increasing Ca^2+^ concentrations under capacitating conditions, Ca^2+^-binding to EFCAB9 is likely to cause changes in the structural conformation of EFCAB9, which affects the interaction with the CatSper channel to control channel activity. The CatSper channel strongly responds to intracellular alkalinization, particularly in the presence of cytosolic free Ca^2+^ (Figure 6A-F). In the absence of EFCAB9-CatSperζ the CatSper channel compromises Ca^2+^ sensitivity in the pH-dependent channel activation.

### EFCAB9 Couples pH-and Ca^2+^ Sensing

Intracellular pH changes rapidly with sperm in response to the environment; CatSper is activated by intracellular alkalinization in mammals and marine invertebrate (Kirichok et al., 2006; Lishko et al., 2010; Lishko et al., 2011; Miller et al., 2015; Seifert et al., 2015; Strunker et al., 2011). However, this pH-dependent activation of CatSper has been explained through speculation about amino acid properties and sequence homology. For example, the remarkably histidine-rich amino terminus of CatSper1 subunit has been proposed to sense intracellular pH or bind to Zn^2+^ (Chung et al., 2011; Kirichok et al., 2006). The inability to heterologously express functional CatSper channels has prevented mutagenesis studies from directly testing these ideas. Interestingly, mouse and human CatSper have differential pH sensitivity despite having similar histidine-rich CatSper1 N-termini (Miller et al., 2015) and pH changes alone are not enough to gate human CatSper channel (Lishko et al., 2010; Strunker et al., 2011). Indeed, intracellular alkalinization is sufficient to activate sea urchin CatSper channels, despite their lack of a histidine-enriched N-terminus (Seifert et al., 2015). These results suggest that CatSper pH-sensing is likely to utilize an evolutionarily conserved mechanism.

We hypothesize that, when closed at rest, CatSperζ-complexed EFCAB9 lies near the cytoplasmic mouth of the pore, inhibiting the channel gating by ensuring its closed conformation. pHi elevation during capacitation partially dissociates EFCAB9 from CatSperζ, which releases gate inhibition and opens the pore, enabling EFCAB9 to bind entering Ca^2+^ and undergo a conformational change to maintain the prolonged open state of the channel. Mammalian EFCAB9 couple pH and Ca^2+^ sensing presumably by adopting several substitutions of the functionally important residues in the EF-hands. For example, glutamate at EF-loop position 12 is preserved in all three EF-hands of sea urchin EFCAB9. However, it is replaced by asparagine in the EFCAB9’s non-canonical EF2 in mammals. The same position in the EFCAB9 EF1 is glutamine, while in human EFCAB9 it is lysine. These changes can contribute to differences in the pH sensitivities between mouse and human CatSper. Likewise, single amino acid substitution of another sperm ion channel also alters the sensitivities to both pH and Ca^2+^. A natural variant (C382R) of SLO3 K^+^ channels of human sperm, which is located between the pore and the cytoplasmic gating ring, has enhanced sensitivities to pH and Ca^2+^(Geng et al., 2017). Thus, loss of negative charges in EFCAB9 should lead to a reduction in binding to Ca^2+^ but might alloW EFCAB9 to acquire pH sensitivity and presumably the interaction with CatSperζ

The absence of EFCAB9 and/or CatSperζ did not completely eliminate the pH dependence nor the requirement of cytosolic Ca^2+^ for CatSper activation (Figures 6 and S6). Such absence, however, makes the complex much less responsive to alkalinization in the presence of physiologically relevant calcium, which ultimately makes the channel less efficient Ca^2+^ conductor. It is possible that additional molecular determinants exist for CatSper pH and Ca^2+^ sensing in mammals. It is intriguing that one of the proteins from the proteomic screen contains C2 Ca^2+^ binding domains, leaving room for future research. Interestingly, amino acid sequences of CatSperζ are variable among mammals along with the histidine-enriched CatSper1-N termini. In contrast, EFCAB9 and the cytoplasmic domains of other pore-forming subunits are evolutionarily conserved (Figures 7C and S6G). It is reasonable to hypothesize that CatSperζ and CatSper1-N termini co-evolved to fine tune variable CatSper pH sensitivity among different mammalian species. Another potential region is CatSper2’s C-terminus. mouse CatSper2 has a ~35 amino acid aspartate-rich region while human CatSper2 has an ~80 amino acid serine-rich region. The species-specific features also provide an explanation for the remaining pH sensor components and the different pH sensitivity between mouse and human CatSper channels.

### Role of EFCAB9-CatSperζ in CatSper Domain Organization

The two-row structure within a single CatSper quadrant is reminiscent of the previously reported flagellar zipper (Friend and Fawcett, 1974; Selvaraj et al., 2007). Whether the CatSper linear domains are the molecular basis of the zipper structure remains to be further explored. Intriguingly, a high-order zipper structure was recently modeled from a quaternary structure of CatSper tetramer, based on the potential cross-linking ability of the evolutionarily conserved cysteine residues of mammalian CatSper α subunits (Bystroff, 2018). Heterologous reconstitution of the CatSper channel for structural studies remains at a standstill. Therefore a possibility of a regularly repeating, quaternary structure of the CatSper channel complex suggests that a direct application of rapidly advancing cryo-EM and cryo-focused ion beam techniques to the sperm tail might be a viable, alternative route for determining molecular details and structure of the CatSper channel. The CatSper complex has multiple transmembrane auxiliary subunits (β, γ δ, and ε) with large extracellular domains. Any information of the sperm surface nanostructure of CatSper will provide a better understanding of these essential, but otherwise unknown, proteins in functional regulation of CatSper.

Our data illustrate that the loss of EFCAB9-CatSperζ results in reduction of the two-row structure to an irregular single structure. While we cannot separate the individual CatSperζ function from EFCAB9, it is reasonable to hypothesize that CatSperζ serves a role in membrane trafficking and scaffolding. CatSperζ might adapt to cytoskeletal structures for the CatSper complex to traffic to the flagellar membrane domains. The fibrous sheath, a cytoskeletal structure unique to the mammalian sperm, lies under the flagellar membrane of the principal piece (Eddy, 2007. Thus, mammalian-specific CatSperζ (Chung et al., 2017) may support the linear arrangement of the CatSper channel complex by linking to an FS component during flagellar development. At the same time, CatSperζ interacts with the pore-forming CatSper α subunits and EFCAB9 (Figures 4, 7, and S6), thus bringing EFCAB9 close to the pore. The CatSperα subunits may form a tetrameric pore via coiled-coil domains in the C-terminus (Lobley et al., 2003), leaving the N-terminal domains as good candidate EFCAB9-CatSperζ interaction site(s). Surprisingly, neither the purified CatSper1 N- nor C-terminal domain directly bind the EFCAB9-CatSperζ complex or CatSperζ alone. These results suggest that the interaction of EFCAB9-CatSperζ with the pore is not binary between CatSper1 and CatSperζ, but rather complex, probably involving multiple pore subunits. Future studies should further enlighten the molecular mechanisms governing the coupling between the CatSper domain organization and channel activity. Other proteins identified in the comparative proteome screen will serve as a foundation to this end.

In summary, we have gained fundamental insights on the regulatory mechanisms of the CatSper channel activity and its domain organization. First, we have identified EFCAB9 as an evolutionarily conserved component of the CatSper channel and provided evidence that the EFCAB9-CatSperζ binary complexes are master integrators linking pH and Ca^2+^ sensing. CatSperζ-complexed EFCAB9 normally limits Ca^2+^ gating prior to alkalinization but, when pH_i_,rises, CatSperζ-free EFCAB9 confers pH-dependent activation of CatSper channels. This coordination is achieved by the Ca^2+^ sensitive and pH-dependent interaction of EFCAB9 and CatSperζ. we further show that EFCAB9-CatSperζ is associated with the channel pore and required for the two-row structure of each single CatSper linear domain, providing a precedent for a linking mechanism for organizing suprastructural domain organization. This study highlights the dual function of EFCAB9-CatSperζ in gatekeeping and domain organization of CatSper.

## EXPERIMENTAL MODEL AND SUBJECT DETAILS

### Animals

*CatSper1, d*, and z-null mice generated in the previous studies (Chung et al., 2017; Chung et al., 2011; Ren et al., 2001) are maintained on a C57/BL6 background. Four week old B6D2F1 females for *in vitro* fertilization (IVF) and 7 week old wt male and female C57BL/6 mice for breeding were purchased from Charles River laboratories. Mice were treated in accordance with guidelines approved by the Yale Animal Care and Use Committees and Boston Children’s Hospital (IACUC).

#### Generation of EFCAB9-null mice by CRISPR/Cas9 and genotyping of mutation

EFCAB9^-^null mice were generated on a C57BL/6 background using CRISPR/Cas9 system. Female mice were superovulated and mated with males to obtain fertilized eggs. pX330 plasmid expressing guide RNA (5’-CCGCCATGAAACTGACTCCGGGG-3’) targeting the first exon of mouse *EFCAB9* was injected into the pronuclei of the fertilized eggs. The developing 2-cell embryos were transplanted into pseudopregnant females. The target region was PCR amplified from alkaline-lysed tail biopsies from founders, and the resulting PCR fragments were subjected to surveyor analysis to examine CRISPR/Cas9 edited indels on the *EFCAB9* locus. Mono-allelic founders with 5bp or 28bp deletion in the first exon were backcrossed with wt C57/BL6 animals to test germline transmission of the mutant alleles. After the mutant *EFCAB9* lines were established, genotyping was performed by multiplex qPCR *(EFCAB9,* forward: 5’-GAAAGCTGCCGCCATGAA-3’, reverse: 5’-ACAGTAAGCAGTAGTTTTGTCCAT-3’, and Probe: 5’-(HEX)-CACAGAAAACACCCCGGAGT CAGT-3’; Internal control (IC), forward: 5’-CACGTGGGCTCCAGCATT-3’, reverse: 5’-TCACCAGTCATTTCTGCCTTTG-3’, and Probe: 5’-(TEX61 5)-CCAATGGT CGGGCACT GCT CAA-3’, BioRad). Potential off-target regions were selected from E-CRISP. Genomic DNA from wt and *EFCAB9-null* mice was extracted. Genomic region containing expected off-target sites were amplified and PCR products were sequenced.

#### Generation of EFCAB9;CatSperz-null mice

*EFCAB9-/-; CatSperz-/-* double knockout mice were generated by mating *EFCAB9+/-* males and *CatSperz-/-* females. Double knockout mice were obtained by mating between double heterozygous *(EFCAB9+/-; CatSperz +/-*) males and females. Genotyping was performed by multiplex qPCR (*CatSperz*, forward. 5’-GCCCAT CTACACCAACGTAACC-3’, reverse. 5’-AGTAACAACCCGTCGGATTCTC-3’, and Probe: 5’-(Cy5.5)-CGGTCAATCCGCCGTTTGTTC-3’; EFCAB9 and IC primers information as stated for *EFCAB9*, BioRad).

### Cell Lines

#### Mammalian cell lines

HEK293T cells (ATTC) were cultured in DMEM (Gibco) containing 10% FBS (Thermofisher) and 1× Pen/Strep (Gibco) at 37 °C, 5% CO2 condition. HEK cells stably expressing human CatSper1 tagged with GFP at C-terminus (HEK-hCatSper1-GFP, a kind gift from David Clapham, Janelia/HHMI) were cultured in 1:1 mixture of DMEM and Ham F12 (DMEM/F12) supplemented with 10% FBS, 1 × Pen/Strep, and 500 μg/ml concentration of geneticin (G418) (Gibco).

#### Bacterial strains

NEB10β (NEB) and BL21-CodonPlus(DE3)-RIL (Agnent Technologies) bacterial strains were used for the molecular cloning and recombinant protein expression, respectively. To express recombinant proteins, freshly formed colonies were picked and inoculated to Luria broth (Sigma-Aldrich) supplemented with 50 μg/m l of chloramphenicol (AmericanBio Inc) and 100 μg/ml of ampicillin (AmericanBio Inc) or 50 μg/m l of kanamycin (AmericanBio Inc) depending on the antibiotic-resistant genes encoded by transformed plasmids. After overnight culture at 37 °C, saturated cultivates were inoculated to terrific broth without antibiotics in one to fiftieth ratio (v/v). Protein expression was induced by Isopropy|-1-thio-β-D-galactopyranoside (IPTG) (AmericanBio Inc) when absorbance value of the culture (OD_600_) reached 0.5-0.7. After IPTG induction, cells were cultured at 16 °C for 14-16hr and harvested to extract recombinant proteins.

### Mouse Sperm Preparation and *In Vitro* Capacitation

Epididymal spermatozoa were collected by swim-out from caudal epididymis in M2 medium (EMD Millipore). Collected sperm were incubated in human tubular fluid (HTF) medium (EMD Millipore) at 2 x 10^6^ cells/ml concentration to induce capacitation at 37 °C, 5% CO_2_ condition for 90 min.

### Human Sperm Preparation

Frozen vials of human sperm from healthy, normal donors were purchased (Fairfax Cryobank). Vials were thawed and mixed with pre-warmed HEPES-buffered saline (HS) (Chung et al., 2017), followed by washing in HS two times. Washed sperm were placed on top of 20% Percoll (Sigma Aldrich) in HS and incubated at 37 °C for 30 minutes to allow for motile sperm to swim-into the Percoll layer. After removing the top layer containing immotile fraction, sperm cells with high motility were collected by centrifugation at 2,000 x g and resuspended in HS.

## METHOD DETAILS

### Antibodies and Reagents

Rabbit polyclonal antibodies specific to mouse CatSper1 (Ren et al., 2001), 2 (Quill et al., 2001), 3, 4 (Qi et al., 2007), β, δ (Chung et al., 2011 Chung et al., 2017 were described previously. To produce EFCAB9 antibody recognizing both mouse and human EFCAB9, peptide corresponding to mouse EFCAB9 (137-154, KKQELRDLFHDFDITGDR) was synthesized and conjugated to KLH carrier protein (Open Biosystems). Antisera from the immunized rabbits were affinity-purified using the peptide immobilized Amino Link Plus resin (Pierce). All the other antibodies and reagents used in this study are commercially available and listed in the key resource table. All the chemicals were from Sigma Aldrich unless indicated.

### Proteomics Analysis

Whole sperm proteome analysis from wt and *CatSper1*-null mice was performed concomitantly with the previous phosphotyrosine proteome analysis (Chung et al., 2014). In short, sperm cells from wt and *Cat Sp er1*-null mice (N=3 per each group) incubated under capacitating conditions were lysed with urea in triplicate. The lysates were reduced, alkylated, trypsin-digested, and reverse-phase purified (Villén and Gygi, 2008). Digested peptides were labeled with TMT to quantify the protein from wt and *CatSper1*-null sperm. The labeled peptides were subjected to mass spectrometric analysis using an Orbitrap Fusion mass spectrometer (Thermo Scientific) with anMS^3^ method. Tandem mass spectrometry (MS/MS) spectra was matched to peptide using Sequest search engine (Thermo Scientific). Peptides having a total of the TMT reporter ion signal/noise ≥ 65 were quantified. False discovery rate was controlled to 1% at the peptide and protein level.

### Open Database Search

#### Transcriptome database

Gene expression data of *EFCAB9, Als2cr11, Slco6c1, Trim69*, and *Fancm* in mouse tissues were obtained from Mouse ENCODE transcriptome data (Yue et al., 2014) as curated in NCBI Gene database.

#### Genome database

Orthologues of CatSper subunits, EFCAB9, ALS2CR11, SLCO6C1, and TRIM69 proteins in 21 eukaryotes were searched in NCBI gene database or by sequence homology analysis as previously described (Chung et al., 2017). Orthologues not annotated in NCBI gene database were identified by comparing amino acid sequences of human orthologue using BlastP in NCBI (http://blat.ncbi.nlm.nih.gov) or JGI genome portal (http.//genome.jgi.doe.gov) with default option. Resulting hits with expected values <10 ^−10^ from Blast alignment were considered as orthologues of the query proteins.

### Multiple Tissue RT-PCR

PCR was carried out using a commercial multiple cDNA panel (MTC, Clontech). Primer pairs amplifying *EFCAB9* (forward. 5’-GAAACGTGAAAGCCTTGATGG-3’ and reverse. 5’-AT CCCGAT CT GT GACTT GTT C-3’), *Efcab1* (forward: 5’-GGACAGAGTATTT CGAGGCTTT-3’ and reverse: 5’-GCCATCACCATTCAGATCAAAC-3’), and *Cam1* (forward: 5’-CAACGAAGTGGATGCTGATG-3’ and reverse. 5’-GCACTGATGTAACCATTCCCA-3’) mRNA were used to examine tissue expression of each gene. *Gapdh* (forward. 5’-TGAAGGTCGGTGTGAACGGATTTGGC-3’ and reverse. 5’-ATGTAGGCCATGAGGTCCACCAC-3’) was used for a control.

### RNA Extraction, cDNA Synthesis, and Real-Time PCR

Total RNA was extracted from testes of 7, 14, 21, 23, 49, and 80 days old wt males using RNeasy mini-kit (Qiagen). 500 ng of RNA was used to synthesize cDNAs using iScript cDNA Synthesis (BioRad). cDNAs were used for real-time PCR (CFX96, Biorad) using the primer pairs. *CatSper1* (forward: 5’-CTGCCTCTTCCTCTTCTCTG-3’ and reverse: 5’-TGTCTATGTAGATGAGGGACCA-3’), *CatSperz* (forward: 5’-GAGACCTCCTTAGCATCGTC-3’ and reverse. 5’-TCGTGGACCTATATGTGATGAG-3’), and *18srRNA* (forward: 5’-CGTCTGCCCTATCAACTTTC-3’ and reverse. 5’-GTTTCT CAGGCTCCCTCTCC-3’). Primers described in *Multiple Tissue Expression* were used to amplify *EFCAB9*, *Efcab1*, and *Calm1* mRNA. *18srRNA* was used as a reference gene to normalize quantitative expression by ddCt method. Three sets of experiments were conducted independently.

### Molecular cloning

#### Mammalian cell expression constructs

Mouse *EFCAB9* ORF clone (Dharmacon, clone 6771929) was subcloned into phCMV3 (*phCMV3-mEfcab*9) to express mouse EFCAB9 tagged with HA at C-terminus. Mouse *CatSperz* (*pCAG-mCatSperz-V5*) (Chung et al., 2017) and *EFCAB9* were subcloned into phCMV3 together for bi-cystronic expression (*phCMV3-mCatSperz-V5-p2A-mEFCAB9*). Human *EFCAB9* ORF (GenScript, #ohu00121d) and *CatSperz* (Chung et al., 2017) were also subcloned into pcDNA3.1 for bi-cystronic expression (pcDNA3.1(-)-*hEFCAB9-HA-p2A-hCatSperz-MYC*). Mouse *CatSper1, 2,* 3, and *4* (Chung et al., 2014) were subcloned into phCMV3 to express C-terminal Flag-tagged proteins (*phCMV3-mCatSper1, 2, 3, or 4-Flag*). Two ORF clones of human *CatSpere* variants (Dharmacon, clone 5269307 and 4823002) were assembled using NEBuilder^®^ HiFi DNA Assembly (NEB) and cloned into phCMV3 (*phCMV3-Flag-hCatSpere*) for the expression construct of human *CatSpere* CDS (NM_001130957.1). Human CatSperg ORF was subcloned into phCMV3 (*phCMV3-Flag-hCatSperg*). Both human *CatSpere* and *CatSperg* constructs were tagged with Flag at the N-termini. For Flag-tagged CatSper constructs in phCMV3 vector, a stop codon was placed at the upstream of HA sequences.

#### Bacterial expression constructs

For N-terminal GST-tagged EFCAB9 and 6xHis-tagged CatSperζ expression in bacteria, mouse *EFCAB9* and *CatSperz* cDNAs were subcloned into pGEX-6P2 (*pGEX-6P2-mEfcab9*) or pET43.1 a(+) (*pET43.1a(+)-mEFCAB9*), and pET32a(+) (*pET32a(+)-gb1-mCatSperz*), respectively. To generate constructs expressing EFCAB9 with two (EFCAB9^Mut2^, D72N/E160Q) and four (EFCAB9^4Mut^, D72N/D82N and D149N/E160Q) amino acid mutations on EF-hand loops, sited-directed mutagenized EFCAB9 cDNAs were subcloned into pGEX-6P2 (*pGex-6P2-mEFCAB9^2Mut^* and *pGex-6P2-mEFCAB9^4Mut^*, respectively) using NEBuilder^®^ HiFi DNA Assembly. The construct to express GST-fused N-terminal intracellular domain of CatSper1 (1-150, *pGEX-2T-mCatSper1-N15*) was a kind gift from D.E. Clapham, Janelia/HHMI. For expression of C-terminal intracellular domain of CatSper1, *pET32 (+)-mCatSper1-C-Flag* was constructed by amplifying a fragment of *CatSper1* cDNA encoding amino acids 574-686 tagged with Flag at C-terminus.

### Recombinant protein expression in mammalian cells

HEK 293T cells were transiently transfected with plasmids encoding various CatSper subunits in combination. the pore-forming (mouse CatSper 1, 2, 3, and 4) or the auxiliary (mouse CatSperβ, δ,ζ, EFCAB9, human CatSperγ, ε, EFCAB9, or mouse and human CatSperζ and EFCAB9 bi-cystronically) subunits. HEK cells stably expressing GFP-tagged human CatSper1 at C-terminus were transfected with a plasmid encoding human CatSperζ and human EFCAB9 bi-cistronically. Lipofectamin 2000 (Invitrogen) or polyethyleneimine (PEI) reagent were used for transfection following the manufacturer’s instruction. Transfected cells were used to characterize antibodies or co-immunoprecipitation (co-IP) experiments.

### Protein Extraction, Immunoprecipitation, and Western blotting

Mouse and human sperm protein was extracted as previously described (Chung et al., 2017, Chung et al., 2011; Chung et al., 2014). Transfected HEK293T cells and HEK cells expressing GFP-tagged human CatSper1 were lysed with 1% Triton X-100 in PBS containing EDTA-free protease inhibitor cocktail (Roche) by rocking at 4 °C for 1 hr, and centrifuged at 14,000 x g for 30 minutes at 4 °C. Solubilized proteins in the supernatant were mixed with Protein A/G-magnetic beads conjugated with either 1 μg each of rabbit polyclonal antibodies (GFP (FL, SantaCruz), mouse CatSper1, mouse CatSperδ, mouse CatSperζ, human CatSperζ) or mouse monoclonal antibodies (Flag (clone M2, Sigma-Aldrich), MYC (9E10, SantaCruz), HA magnetic beads (clone 2-2.2.14, Pierce)) or V5 agarose (clone V5-10, Sigma-Aldrich). The immune complexes were incubated at 4 °C overnight and co-IP products were eluted with 60 μl of 2x LDS sampling buffer supplemented with 50 μM dithiothreitol (DTT) and denatured at 75 °C for 10 minutes. Primary antibodies used for the western blotting were: rabbit polyclonal anti-mouse CatSper1, CatSper2, CatSper3, CatSper4, CatSperβ, CatSperδ, EFCAB9, human CatSperζ GFP (FL, SantaCruz), CaM, (05-173, Upstate), phosphotyrosine (clone 4G10, EMD Millipore) at 1 μg/ml, mouse CatSperζ (2.7 μg/ ml), mouse monoclonal anti-HA (clone 2-2.2.14, Pierce), Flag (clone M2, Sigma-Aldrich) at 0.5 μg/ml, acetylated tubulin (T7451, Sigma Aldrich, 1:20,000), HRP-conjugated V5 (clone E10/V4RR, ThermoFisher, 1:2,000), HA (clone 6E2, CST, 1:1,000), and MYC (Clone 9B11, CST, 1:1,000). For secondary antibodies, anti-mouse IgG-HRP, antirabbit IgG-HRP (Jackson ImmunoResearch, 1:10,000) and mouse IgG Trueblot (clone eB144, Rockland, 1:1,000) were used.

### Sperm Immunocytochemistry

Mouse and human sperm were washed in PBS twice, attached on the glass coverslips, and fixed with 4% para fo rmaldehyde (PFA) in PBS at room temperature (RT) for 10 minutes (mouse) or at 4 °C for 1 hr (human). Fixed samples were permeabilized using 0.1% Triton X-100 in PBS at RT for 10 minutes, washed in PBS, and blocked with 10% goat serum in PBS at RT for 1 hr. Cells were stained with anti-mouse EFCAB9 (20 μg/ml), CatSper1 (10μg/ml), or CatSperζ (20μg/ml) in PBS supplemented with 10% goat serum at 4 °C overnight. After washing in PBS, the samples were incubated with goat anti-rabbit Alexa647 or Alexa555-plus (Invitrogen, 1:1,000) in 10% goat serum in PBS at RT fo r 1 hr. Immunostained samples were mounted with Prolong gold (Invitrogen) and cured for 24 hr, followed by imaging with Zeiss LSM710 Elyra P1 using Plan-Apochro m bat 63X/1.40 and alpha Plan-APO 100X/1.46 oil objective lens (Carl Zeiss). Hoechst dye was used to counterstain nucleus for sperm head.

### Super-Resolution Imaging

#### Structured illumination microscopy (SiM)

Structured illumination microscopy (SIM) imaging was performed with Zeiss LSM710 Elyra P1 using alpha Plan-APO 100X/1.46 oil objective lens. Samples were prepared as described in *Sperm Immunocytochemistry* with a minor modification in that the immunostained sperm coverslips were mounted with VectaSheild (Vector laboratory). 2D SIM images were taken using a laser at 642 nm (150 mW) for Alexa647 (Invitrogen). The images were acquired using 5 grid rotations with a 51 nm SIM grating period. For 3D SIM images, a laser at 561 nm (200 mW) was used for Alexa555-plus (Invitrogen). Z-stack was acquired from 42 optical sections with a 100 nm interval. Each section was imaged using 5 rotations with a 51 nm grating period. Both 2D and 3D SIM Images were rendered using Zen 2012 SP2 software.

#### 4Pi Single-Molecule Switching Nanoscopy (4Pi-SMSN)

Mouse sperm were attached onto the center of 25-mm diameter of glass cover slips. The samples were prepared as described in *Sperm immunocytochemistry* with Alexa647-conjugated 2^nd^ antibody. The samples were imaged with a custom-built 4Pi Single-Molecule Switching Nanoscopy (4Pi-SMSN) system as previously described (Huang et al., 2016) with minor modifications. Briefly, the fluorescent signal was collected coherently by two opposing objectives (100 x/1.35NA, silicone oil immersion, Olympus) and imaged on a sCMOS camera (ORCA-Flash 4.0v2, Hamamatsu). The microscope was equipped with an excitation laser at 642 nm (MPB Communications, 2RU-VFL-2000-642-B1R) and an activation laser at 405 nm (Coherent OBIS 405 LX, 50 mW). All data were acquired at 100 fps at a 642 nm laser intensity of about 7.5 kW/ cm^2^. The full system design and the image analysis algorithms were previously described in detail (Huang et al., 2016). The 4Pi movies were rendered using Vutara SRX software (Bruker). Angular profiling was carried out as previously reported (Chung et al., 2014).

### Scanning Electron Microscopy

Sperm from wt, *CatSper1-null,* and *EFCAB9-null* males were washed in PBS and attached on the glass coverslips. The coverslips were fixed with 4% paraformaldehyde in PBS for 10 minutes at RT, and washed in PBS twice. The coverslips were incubated with 2.5% glutaraldehyde in 0.1M sodium cacodylate buffer pH7.4 for another hour at 4 °C. Samples were then rinsed, post fixed in 2% osmium tetroxide in 0.1 M sodium cacodylate buffer pH 7.4, and dehydrated through a series of ethanol to 100%. The samples were dried using a Leica 300 critical point dryer with liquid carbon dioxide as transitional fluid. The coverslips were glued to aluminum stubs, and sputter coated with 5 nm platinum using a Cressington 208HR (Ted Pella) rotary sputter coater.

### Mating Test and *In Vitro*F ertilizatio

Female mice were caged with heterozygous or homozygous *EFCAB9*- mutant or *EFCAB9*, *CatSperz* double mutant males for two months to record pregnancy and litter size when gave births. For IVF, 5-7 weeks B6D2F1 female mice were superovulated by injecting progesterone and anti-inhibin serum (Central Research Co, Ltd) (Hasegawa et al., 2016), and the oocytes were collected after injecting 13 hr from 5 U of human chorionic gonadotrophin (EMD millipore). Prepared epididymal sperm were capacitated at 37 °C for 90 min, and inseminated to oocytes with 2 x 10^5^ cells/ml concentration. After 5 hr co-incubation, oocytes were washed and transferred to fresh HTF medium, and cultured overnight at 37 °C under 5% CO_2_. 2-cell embryos were counted 20-22 h after insemination as successful fertilization.

### Flagella Waveform Analysis

To tether sperm head for planar beating, non-capacitated or capacitated spermatozoa (2 x 10^5^ cells) were transferred to the fibronectin-coated 37 °C chamber for Delta T culture dish controller (Bioptechs) filled with HEPES-buffered HTF medium (H-HTF) (Chung et al., 2017) for 1 minute. Flagellar movements of the tethered sperm were recorded for 2 sec with 200 fps using pco.edge sCMOS camera equipped in Axio observer Z1 microscope (Carl Zeiss). All movies were taken at 37 °C within 10 minutes after transferring sperm to the imaging dish. FIJI software (Schindelin et al., 2012) was used to measure beating frequency and α-angle of sperm tail, and to generate overlaid images to trace waveform of sperm flagella as previously described (Chung et al., 2017).

### Recombinant Protein Purification

Each Construct expressing GST-tagged EFCAB9, EFCAB9^2Mut^, EFCAB9^4Mut^, and N-terminus of CatSper1 (CatSper-N1 50) and 6xHis-tagged CatSperζ and C-terminus of CatSper1 (CatSper1-C) was transformed to BL21-CodonPlus(DE3)-RIL competent cells (Agilent Technologies). Fresh colonies were inoculated and cultured into LB with antibiotics at 37 °C overnight. Saturated cultivates were 50 times diluted in TB medium and cultured further at 37 °C until the OD_600_ values reach 0.5 - 0.7. To induce protein expression, IPTG (0.2 mM for EFCAB9 and CatSper1-N150; 0.1 mM for CatSperζ; 0.05 mM for EFCAB9^2Mut^, EFCAB9^4Mut^, and CatSper1-C)was added to the bacteria cultures and incubated further for 16 hr at 16 °C. When the CatSper1-N150 recombinant protein expression is induced by IPTG, proteasome inhibitor MG-132 (Calbiochem) idded to 10 μM. Cultured cells were harvested and washed with cold PBS. Cell pellets were was adresuspended in buffers containing 10 mM HEPES in pH 8.0 (EFCAB9, EFCAB9^2^ EFCAB9^4Mut^), pH 7.5 (CatSper1-N150 and CatSper1-C) or pH6.0 (CatSperζ) with 135 mM NaCl containing EDTA-free protease inhibitor cocktail (Roche). MG-132 was also added to 30 μM in the resuspension buffer for CatSper1-N150 recombinant protein purification. Resuspended cells were lysed using Emulsi Fiex-C3 (AVESTIN, Inc.) or VCX500 sonicator (SONICS). Lysates were centrifuged at 14,000 x g for 1 hr at 4 °C. The supernatant were incubated with glutathione agarose (Pierce) or HisPur Ni-NTA resin (Pierce) depending on the tag of the target recombinant proteins for 1 hr at RT. Glutathione agarose was washed with 10 mM HEPES buffer pH7.5, 140 mM NaCl and the GST-fused protein was eluted in the elution buffer (50 mM HEPES buffer pH7.4 with 10 mM reduced glutathione). Ni-NTA resin was washed with 10 mM HEPES buffer pH6.5, 300 mM NaCl with 30 mM (CatSper1-C) or 100 mM (CatSperζ) Imidazole and the His-tagged proteins were eluted with the buffer containing 10 mM HEPES buffer pH7.4, 135 mM NaCl, 300 mM Imidazole. The eluents were dialyzed at 4 °C against storage buffer (10mM HEPES buffer pH7.4, 135mM NaCl in 50% glycerol). To express and purify recombinant EFCAB9 and CatSperζ proteins together, competent cells were transformed *by pET43.1 (+)-mEFCAB9* and *pET32 (+)-mCatSperz* together. A fresh colony was picked, and cultured as described above. To induce the protein expression, the cells were treated with 0.05 mM IPTG and harvested after 14 hr culture at 16 °C. After washing in PBS, the pellet was resuspended in 10 mM HEPES buffer pH7.5, 150 mM NaCl, and lysed using EmulsiFlex-C3 (AVESTIN, Inc.). Lysates were centrifuged and the collected supernatant was incubated with glutathione agarose. The agarose was washed with 10 mM HEPES buffer pH7.5, 150 mM NaCl. The bound recombinant proteins were eluted from the glutathione agarose with the elution buffer for GST-tagged protein, followed by dialysis against the storage buffer as described above.

### Pull-down Assay

Pull-down assay was performed to test direct binding between purified recombinant proteins. 5 μI of glutathione agarose (GST pull-down; Pierce), HisPur Ni-NTA resin (His pull-down; Pierce), or 15 ul of Protein-A/G agarose beads slurry (SantaCruz) cross-linked with CatSper1 antibody was equilibrated with pre-binding buffer (10 mM HEPES pH7.5, 140 mM NaCl), and incubated with the recombinant GST-tagged EFCAB9 or 6xHis tagged CatSperζ, or GST-CatSper1-N150 protein (bait proteins), respectively, at 4 °C for overnight. Incubated resin was washed in prebinding buffer three times and equilibrated in the binding buffer in various compositions for each experiment. Prey proteins subjected to interact with bait proteins were incubated with the resin equilibrated with binding buffer for 1 hr at RT. The resin was washed in the binding buffer for each group four times. Resin was collected and the bound proteins were eluted with 2X LDS sampling buffer supplemented with 50 mM DTT, and denatured at 75 °C for 10 minutes. Protein interaction was confirmed by Coomassie blue staining (GeCode™ Blue Safe Protein Satin, ThermoFisher) or western blotting. Detailed experiment procedures and compositions of binding buffers for each experiment are as described below.

*interaction between EFCAB9 and CatSperζ* Recombinant EFCAB9 and CatSperζ proteins were used as bait for glutathione agarose or His-Pur Ni-NTA resin, respectively to test their interaction reciprocally. In His-pull down assay, pre-binding buffer contains 30 mM Imidazole. Each pre-binding buffer used in GST or His pull-down assay was also used as binding buffer.

*Ca^2+^- or pH-dependent interaction between EFCAB9 and CatSperζ* To examine the Ca^2+^- or pH-sensitivity of the interaction, GST pull-down was performed with the recombinant proteins. Recombinant EFCAB9 bound to glutathione resin was equilibrated with binding buffer at pH7.5 with different Ca^2+^ concentration by adding 2 mM CaCl_2_ or 2 mM EGTA or different pH (pH 6.0, 7.5, and 8.5) with nominal 0 Ca^2+^ concentration (no added Ca^2+^). Calcium and pH-dependent interaction within mutated EFCAB9 and CatSperζ recombinant proteins was performed with the same buffers.

*interaction between EFCAB9-CatSperζ complex and CatSper1 intracellular domains*. Co-purified EFCAB9-CatSperζ recombinant proteins were subjected to interact with either N- or C-terminal domain of CatSper1 protein purified as described. Direct interaction of N-terminal domain (CatSper1-N150) with CatSperζ or EFCAB9-CatSperζ complex was tested by pulling down CatSper1-N150 with glutathione agarose or CatSper1-crosslinked A/G agarose slurry beads. GST pull-down was performed to test the interaction between EFCAB9-CatSperζ complex and CatSper1-C.

### Sequence comparison of orthologs

Protein sequences of EFCAB9, CatSperζ, and intracellular domains of CatSper 1, 2, 3, 4 and Cav3 .1 over 100 amino-acid residues were collected from 12 mammals, mouse (*Mus musculus*), rat (*Rattus norvegicus*), hamster (*Cricetulus griseus*), squirrel (*ictidomys tridecemlineatus*), human (*Homo sapiens*), chimpanzee (*Pan troglodytes*), baboon (*Papio anubis*), pig (*Sus scrofa*), cow (*Bos taurus*), dog (*Canis lupus familiaris*), cat (*Felis catus*), and ferret (*Mustela putorius furo*). Orthologue sequences were aligned by MUSCLE alignment algorithm with default option (Edgar, 2004) and pairwise distances between orthologues were calculated. Pairwise distances were represented to heatmaps indicating sequence diversity among mammals. Co-evolution between two orthologues was modeled by plotting the pairwise distances of the orthologues between two species. Correlation coefficient of the plot is considered as co-evolution score, and the slope value of the linear-regression model indicates relative evolutionary rate within orthologues.

### Electrophysiology

Sperm cells were allowed to adhere to glass coverslips. Gigaohm seals were formed at the cytoplasmic droplet of motile sperm cells in standard HEPES saline buffer containing (in mM): 130 NaCl, 20 HEPES, 10 lactic acid, 5 glucose, 5 KCl, 2 CaCl_2_, 1 MgSO_4_, 1 sodium pyruvate, pH 7.4 adjusted with NaOH, 320 mOsm/L (Lishko et al., 2010; Lishko et al., 2011). Transition into whole-cell mode was achieved by applying voltage pulses (400-620 mV, 1 ms) and simultaneous suction. Data were sampled at 10 Hz and filtered at 1 kHz and cells were stimulated every 5 seconds. The divalent-free bath solution (DVF, ~320 mOsm/L) consisted of (in mM): 140 CsMeSO_3_, 40 HEPES, 1 EDTA and pH 7.4 was adjusted with CsOH. pH-dependent *I_CatSper_* from wt and CatSperz-null sperm recorded with pipette solution (~335 mOsm/L) contained (in mM): 130 CsMeSO_3_, 60 MES, 3 EGTA, 2 EDTA, 0.5 Tris-HCl. Access resistance was 43-62 MΩ. To induce intracellular alkalization (pH, = ~7.4), 10 mM NH4Ci were added to the bath solution. For experiments with defined intracellular pH and 0 μM or 10 μM Ca^2+^, inside pipette solution containing (in mM): 130 CsMeSO_3_, 60 MES (pH = 6.0) or HEPES (pH = 7.4), 5 EGTA, 4 CsCl, and 2 EDTA (0 μM Ca^2+^) or 130 mM CsMeSO_3_, 65 MES (pH = 6.0) or 60 HEPES (pH = 7.4), 4 CsCl and 2 HEDTA (10 μM Ca^2+^). Required CaCl_2_ concentrations for desired free Ca^2+^ concentration were calculated with WinMAXC32 version 2.51 (Chris Patton, Stanford University). Access resistance was 42-61 MΩ. pH of pipette solution was adjusted with CsOH. All experiments were performed at 22 °C. D ata were analyzed with Clampfit (v10.3, pClamp) and OriginPro (v9.0, Originlab).

## QUANTIFICATION AND STATISTICAL ANALYSIS

Statistical analyses were performed using Student’s t-test. Differences were considered significant at *\*P*<0.05, \*\**P*<0.01, and \*\*\**P*<0.001. Pairwise distances between orthologues were calculated using MEGA6 software (Tamura et al., 2013). Correlation coefficient and linear regression model in orthologue sequence comparison were calculated using R software.

## DATA AND SOFTWARE AVAILABILITY

The original dataset is available at doi.10.17632/mv52 ykks6x.1

## ACKNOWLEDGMENTS

We thank David E. Clapham for sharing reagents (*EFCAB9*founder mice, hCatSper1-GFP cell line, *pGEX2T2-mCatSper1-N150*) and critical reading of the draft, Sang-Hee Shim for sharing algorithm for angular plots, Jong-Nam Oh for assistance in immunocytochemistry with human sperm samples, and the Yale Center for Cellular and Molecular Imaging for assistance in scanning electron microscopy. This work was supported by start-up funds from Yale University School of Medicine, a Goodman-Gilman Scholar Award-2015, and a Rudolf J. Anderson Fellowship award to J.-J.C., by NIH R01GM111802, Pew Biomedical Scholars and Rose Hill awards, and Packer Wentz Endowment Will to P.V.L., and by the Wellcome Trust (203285/B/16/Z) and the Yale Diabetes Research Center (NIH P30 DK045735) to J.B.

## AUTHOR CONTRIBUTION

J.-J.C. conceived and supervised the project. J.-J.C. and J.Y.H. designed, performed, and analyzed experiments. J.-J.C. created Efcab9-null mice, did initial characterization, and contributed to all of the 4Pi-SMSN and SEM imaging. J.Y.H. performed comparative genomic screens, molecular and cell biology and protein chemistry experiments including expression construct generation, protein purification, immunocytochemistry, confocal and SIM imaging, and motility analysis. N.M. did electrophysiological recordings and N.M., P.V.L., and J.-J.C. analyzed the electrophysiological data. Y.Z. performed 4Pi-SMSN imaging, image analysis, and rendering. J.-J.C. and R.A.E. performed proteomic experiments, and J.-J.C., J.Y.H., and R.A.E. analyzed the proteomic data. S.P.G., J.B., and P.V.L. provided crucial reagents and equipment. J.Y.H. and J.-J.C. assembled figures and wrote the manuscript with the input from the co-authors.

